# Computational and functional characterization of the PI(4,5)P_2_ binding site of the TRPM3 ion channel

**DOI:** 10.1101/2022.06.05.494899

**Authors:** Siyuan Zhao, Vincenzo Carnevale, Eleonora Gianti, Tibor Rohacs

## Abstract

Transient Receptor Potential Melastatin 3 (TRPM3) is a heat-activated ion channel expressed in peripheral sensory neurons and the central nervous system. TRPM3 activity depends on the membrane phospholipid phosphatidylinositol 4,5-bisphosphate [PI(4,5)P_2_], but the molecular mechanism of activation by PI(4,5)P_2_ is not known. As no experimental structure of TRPM3 is available, we built a homology model of the channel in complex with PI(4,5)P_2_ *via* molecular modeling. We identified putative contact residues for PI(4,5)P_2_ in the pre-S1 segment, the S4-S5 linker, and the proximal C-terminal TRP-domain. Mutating these residues increased sensitivity to inhibition of TRPM3 by decreasing PI(4,5)P_2_ levels by phosphatidylinositol 4-kinase inhibition. Changes in ligand-binding affinities *via* MM/GBSA showed reduced PI(4,5)P_2_ affinity for the mutants. Mutating PI(4,5)P_2_ interacting residues also reduced sensitivity for activation by the endogenous ligand pregnenolone sulfate (PregS), pointing to an allosteric interaction between PI(4,5)P_2_ and PregS. Mutating residues in the PI(4,5)P_2_ binding site in TRPM8 had similar effects, increased sensitivity to PI(4,5)P_2_ depletion, and reduced sensitivity to menthol. Mutation of most PI(4,5)P_2_ interacting residues in TRPM3 also increased sensitivity to inhibition by Gβγ, indicating allosteric interaction between Gβγ and PI(4,5)P_2_. Disease-associated gain of function TRPM3 mutations on the other hand, resulted in no change of PI(4,5)P_2_ sensitivity, indicating that mutations did not increase channel activity *via* increasing PI(4,5)P_2_ interactions. Our data provide insight into the mechanism of regulation of TRPM3 by PI(4,5)P_2_, its relationship to endogenous activators and inhibitors of TRPM3, as well as identify similarities and differences between PI(4,5)P_2_ regulation of TRPM3 and TRPM8.

## INTRODUCTION

TRPM3 is a heat-activated, outwardly rectifying, Ca^2+^ permeable non-selective cation channel expressed in a variety of tissues, including peripheral sensory neurons of the Dorsal Root Ganglia (DRG), and the central nervous system (1). Its chemical activators include the endogenous neurosteroid pregnenolone sulfate (PregS) (2) and the synthetic compound CIM0216 (3). TRPM3 activity can be inhibited by a number of compounds, including natural flavonones, such as isosakuranetin (4), the non-steroid anti-inflammatory drug diclofenac, and the anti-epileptic medication primidone (5). TRPM3 is a very well established peripheral noxious heat sensor. Genetic deletion of this channel in mice results in impaired noxious heat sensation (6–8), and impaired inflammatory thermal hyperalgesia (6,8). TRPM3 inhibitors also reduce thermal hyperalgesia and basal heat sensitivity (4,5,8).

Activation of Gi-coupled receptors inhibits TRPM3 activity. This effect was demonstrated both by native receptors in DRG neurons, including μ-opioid and GABA_B_ receptors (9–11), as well as by heterologously expresses Gi-coupled receptors such as M2 muscarinic receptor and μ-opioid receptors (9,10). Inhibition by Gi-coupled receptors is mediated by direct binding of Gβγ to the channel protein (9–11) through a short α-helical peptide encoded by an alternatively spliced exon in TRPM3, the co-structure of which with Gβγ has been recently determined by X-ray crystallography (12). (TRPM3 has a large number of splice variants, and some of the alternatively spliced exons are in the N-terminus (1), which makes residue numbering confusing (13–15).) The Gβγ binding peptide is present in TRPM1, the closest relative of TRPM3, which is also inhibited by Gβγ (16), but it is missing from the rest of the TRPM family. Activation of recombinant (9) or native (17) Gq-coupled receptors may also inhibit TRPM3, which is also mediated mainly by Gβγ binding (9).

It was recently shown that mutations in TRPM3 are associated with developmental and epileptic encephalopathies manifesting as intellectual disability and seizures in children (13). The originally described two disease-associated mutations both showed a gain of function phenotype with increased basal activity and increased heat and agonist sensitivity (14,15). This points to the importance of TRPM3 in the brain, but knowledge on the functional role of TRPM3 in the central nervous system is quite limited (18).

Phosphoinositides, especially PI(4,5)P_2_ are common ion channel regulators (19,20). Most TRP channels, including TRPM3 (21,22) are positively regulated by phosphoinositides (23), but in some cases such as TRPV1 (24,25), or TRPC channels (26,27) this regulation is complex, and sometimes controversial, with both negative and positive effects having been proposed. With the exception of TRPM1, which is very difficult to study in expressions systems, all members of the TRPM subfamily have been shown to be positively regulated by PI(4,5)P_2_ and no negative regulation has been proposed for any TRPM subfamily member (23). While cryogenic electron microscopy (cryo-EM) structures are available for five out of eight members, the only TRPM channel where the PI(4,5)P_2_ binding site is revealed by structural studies, is TRPM8 (28).

Currently, it is not known which residues in the TRPM3 protein PI(4,5)P_2_ binds to, and there is no experimentally determined structure available for TRPM3. To fill this key gap in knowledge, we generated a homology model of TRPM3, based on the experimental structure of TRPM4 in the ligand-free (apo) state (29). We then docked PI(4,5)P_2_ to our model of TRPM3, and identified putative PI(4,5)P_2_ interacting residues in the pre-S1 segment, the S4-S5 linker, and the proximal C-terminal TRP-domain. We validated our results by docking PI(4,5)P_2_ to an apo structure of TRPM8 (30), which showed remarkable similarity to the TRPM8-PI(4,5)P_2_ structures (28), experimentally determined recently. *In silico* mutations of the PI(4,5)P_2_ contact residues in TRPM3, followed by ligand-binding affinity changes via Molecular Mechanics/Generalized Born Surface Area (MM/GBSA), showed reduced PI(4,5)P_2_ binding affinity to TRPM3. We experimentally validated the importance of these residues by demonstrating that their mutations increased sensitivity to inhibition by PI(4,5)P_2_ depletion in electrophysiology experiments. We also showed that mutating most of these residues increased sensitivity to Gβγ inhibition, and decreased sensitivity to agonist activation, indicating allosteric interaction between PI(4,5)P_2_ and endogenous activators and inhibitors. Furthermore, we demonstrated that gain of function disease-associated mutations did not change PI(4,5)P_2_ sensitivity, indicating that the mutations do not increase channel activity *via* promoting PI(4,5)P_2_ activation. Our data provide mechanistic insights into regulation of TRPM3 by its key endogenous cofactor PI(4,5)P_2_.

## RESULTS

Our goal in this study was to identify the PI(4,5)P_2_ binding site of TRPM3. As there is currently no experimentally determined TRPM3 structure available, we generated a homology model of the human TRPM3 based on the cryo-EM structure of the mouse TRPM4 in the apo state (6BCJ) (29) (**Fig. 1**). The template was selected as the closest homolog to TRPM3 in the TRPM family with an experimental structure available when the model was built (see the Methods section, and **Scheme S1** for details). Our model of TRPM3 aligns very well with the models of TRPM3 from different organisms generated recently by AlphaFold (31,32) (**Fig. S1**), as well as with a model of TRPM3 obtained using the experimental structure of mouse TRPM7 (33) in EDTA (5ZX5) as the template (**Fig. S2**), all of which became available after our original homology model was built, providing *a posteriori* validation of our TRPM3 model.

**Figure 1.**
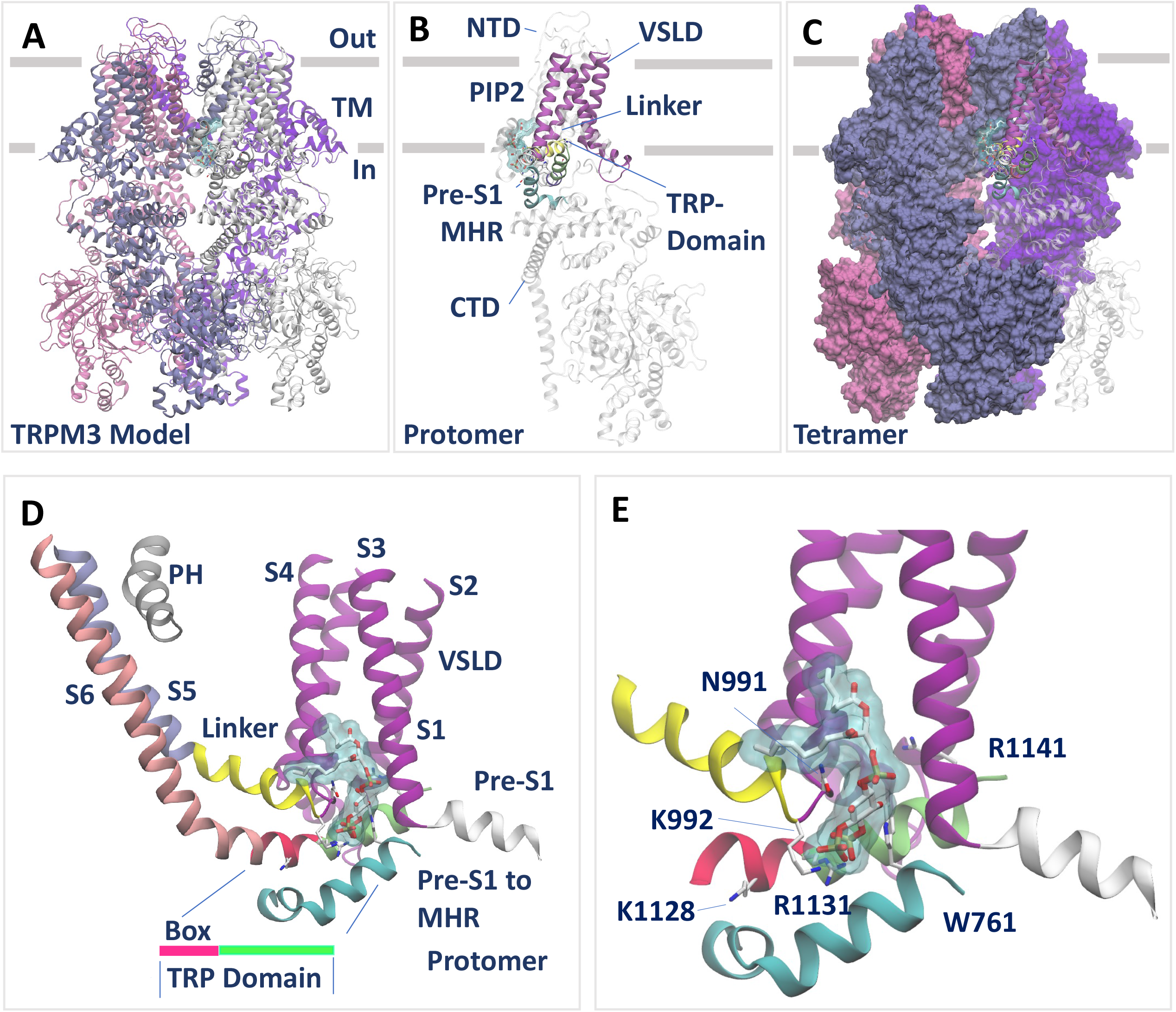
Model of TRPM3 in complex with PI(4,5)P_2_. The homology model of TRPM3 was built based on the structure of TRPM4 (6BCJ) as described in the Results and in the Methods sections. For simplicity, only one molecule of PI(4,5)P_2_ is shown. (**A** to **C**) Views from the transmembrane (TM) plane of TRPM3 tetramer (A and C) and protomer (B). In (A), protein atoms are shown in new cartoon representation, each in a different color. PI(4,5)P_2_ atoms are shown in licorice representation, with C, N, O atoms colored in white, blue and red, respectively. PI(4,5)P_2_ atoms are also shown in surface representation, colored in light blue (transparent). In (B), protein atoms of one protomer are shown in new cartoon representation. Atoms in the pre-S1 domain, voltage-sensor-like domain (VSDL), the S4-S5 linker, the S5, S6 and the pore (PH) helices are shown in new cartoon representation, colored in cyan, magenta, yellow, ice blue, grey and pink, respectively. The TRP domain is colored in magenta (TRP-box) and green. The remaining structural elements in the protomer are shown in white (transparent). PI(4,5)P_2_ is shown as in (A). In (C), protein atoms of three of the four protomers are shown in surface representation, each in a different color. The fourth protomer is represented as in (B). (**D** and **E**) Close-up views of the PI(4,5)P_2_ binding site in TRPM3. The PI(4,5)P_2_ binding site comprises elements from an individual protomer, as highlighted in (C), where no overlap is shown with elements from an adjacent protomer (colored in ice blue). In (E), zoom-in to the binding site residues. Protein atoms are represented as new cartoons. In (D and E), PI(4,5)P_2_ atoms are represented as in (A).

**Scheme S1.**
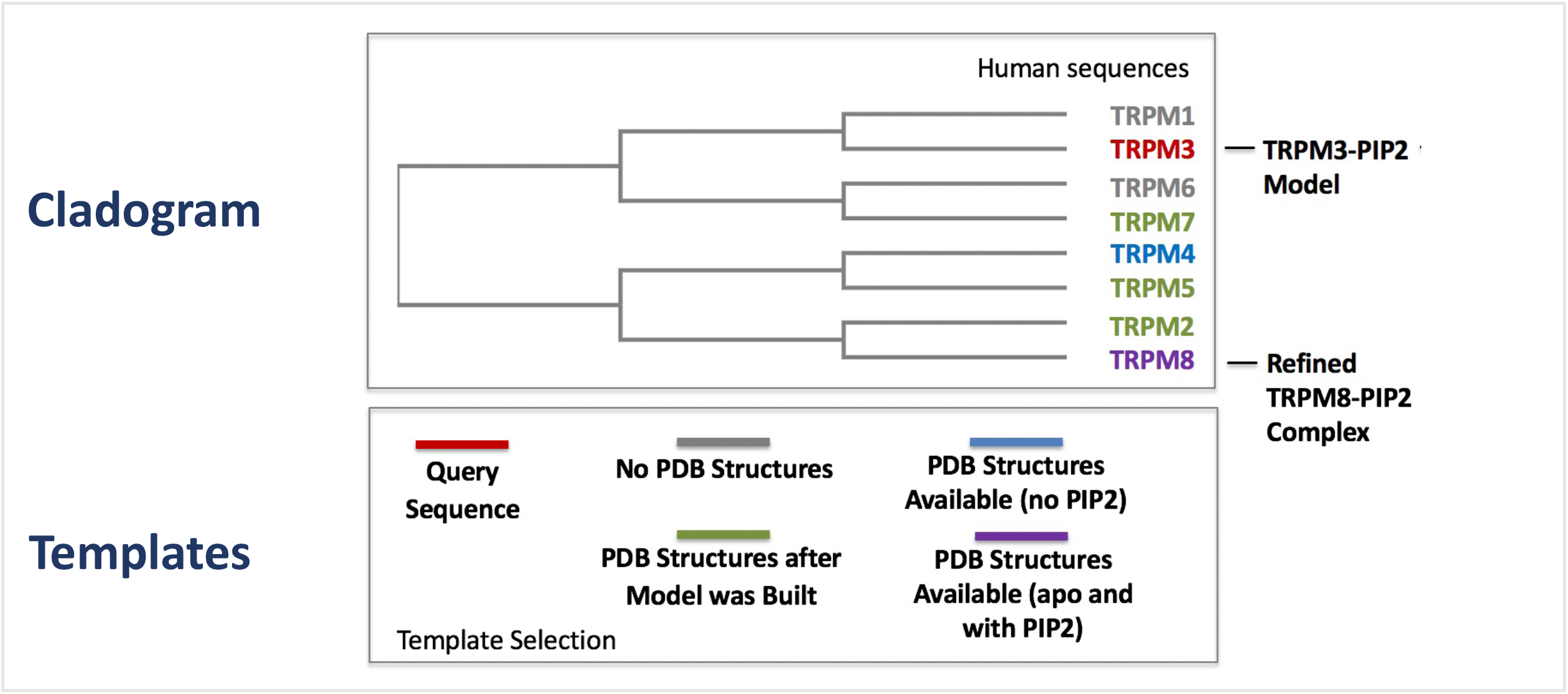
Cladogram and inventory of experimental structures available as templates for building the model of TRPM3 in complex with PI(4,5)P_2_. When the model was built, the structure of TRPM4 in a ligand free state was used as the template (PDB-ID 6BCJ) (29). This structure was selected as the closest homolog to TRPM3 in the TRPM family with an experimental structure available at the time.

Next, we identified putative residues interacting with PI(4,5)P_2_ in TRPM3 by using two complementary approaches. *First*, we scanned the surface of apo TRPM3 (model built on TRPM4) for putative binding sites using the program SiteMap (34,35). *Second*, we relied on sequence and structural information available on TRPM8 to detect, by homology, which residues are likely to interact with PI(4,5)P_2_. Specifically, (I) we generated a sequence alignment of TRP-domain residues, K995, R998 and R1108 in the rat TRPM8, which are conserved among TRPM family members (**Fig. 3A**) and were previously suggested to play a key role in PI(4,5)P_2_ interactions (36); and (II) starting from the apo cryo-EM structure of the flycatcher apo TRPM8 (fcTRPM8) (6BPQ) (30), we built a refined model of this channel bound to PI(4,5)P_2_ at a site comprising the conserved TRP-domain residues (**Fig. 2**). Comparing this complex with the apo-TRPM3 model showed that the most suitable site for lipid binding (i.e. the top-scoring binding spot combining SiteMap predictions and structural information) in TRPM3 corresponded to the PI(4,5)P_2_ site identified in TRPM8. We used this lipid binding site to generate a model of TRPM3 in complex with a version of PI(4,5)P_2_ with truncated tails (similar to the synthetic diC_8_ PI(4,5)P_2_, which is fully functional in experiments) by molecular docking using the program Glide (37). We ranked the lipid binding modes by the standard precision scoring function. The best binding mode, defined as the best docking score (kcal/mol) obtained at the binding site common to TRPM8, is shown in **Figure 1**.

**Figure 2.**
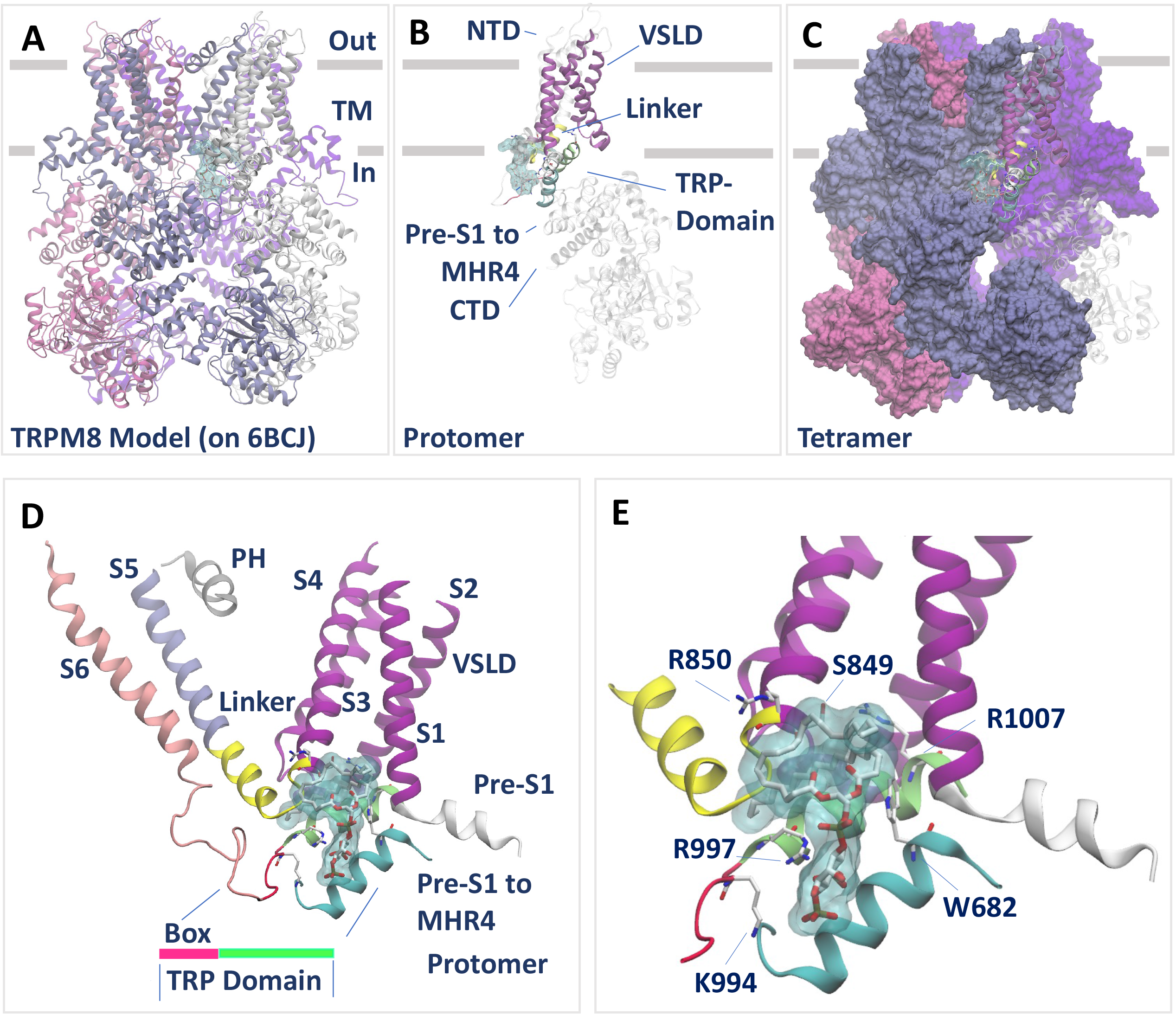
Refined model of TRPM8 in complex with PI(4,5)P_2_. PI(4,5)P_2_ was docked to the apo structure of the fcTRPM8 (6BPQ) (30) as described in the Results and in the Methods sections. For simplicity, only one molecule of PI(4,5)P_2_ is shown. (**A** to **C**) Views from the transmembrane (TM) plane of TRPM8 tetramer (A and C) and protomer (B). (**D** to **E**) Close-up views of the PI(4,5)P_2_ binding site in TRPM8. All representations are reproduced as in Figure 1.

**Figure 3.**
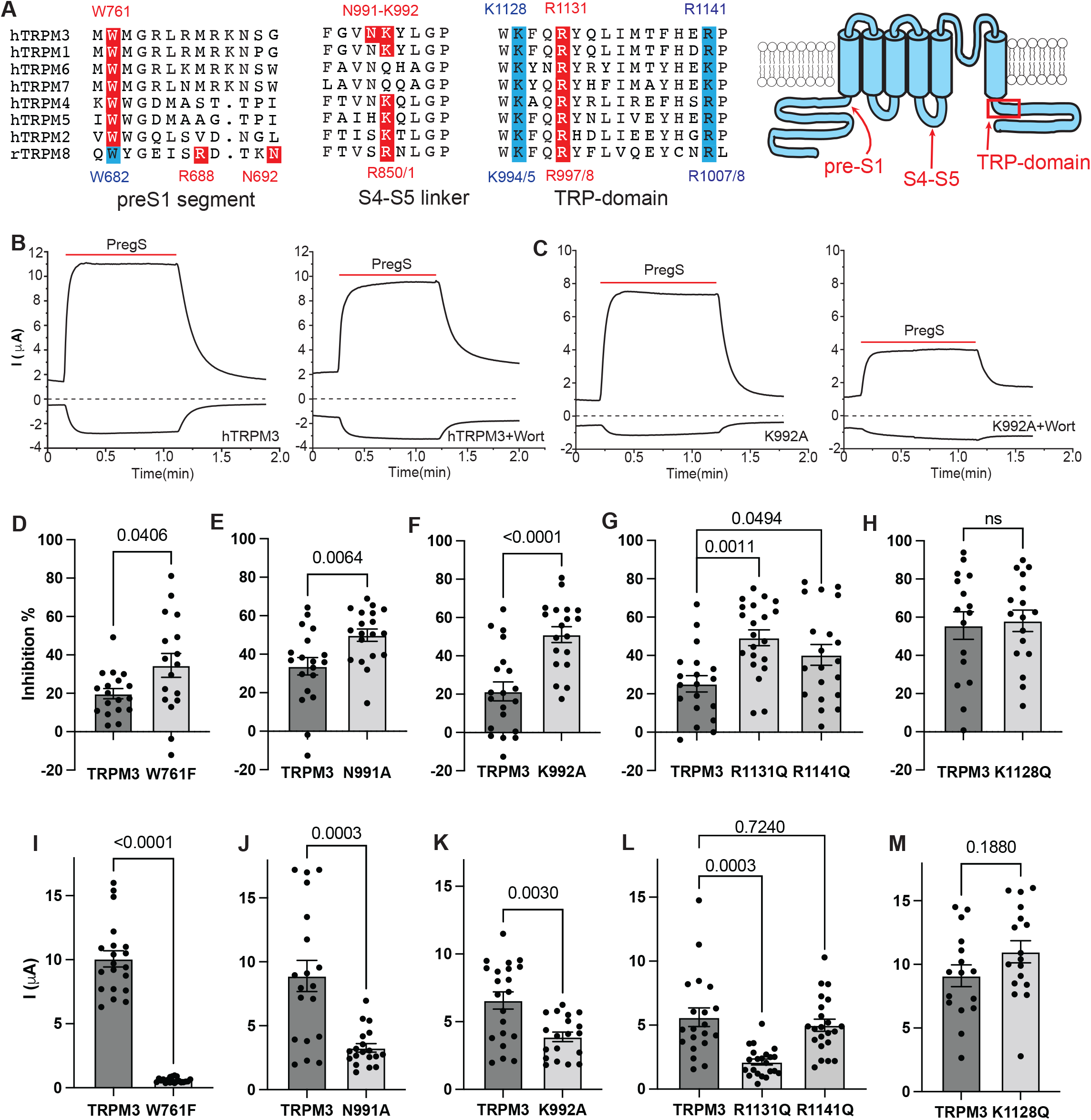
Mutating putative PI(4,5)P_2_ interacting residues increases sensitivity of TRPM3 to inhibition by PI(4,5)P_2_ depletion. (A). Sequence alignment and cartoon of the PI(4,5)P_2_ interacting regions in TRPM3 and TRPM8. Red residues are in contact with PI(4,5)P_2_ in the TRPM8 cryoEM structure(s) and/or in our TRPM3 model, when conserved in other TRPM channels, they are also labeled red. Residues in cyan in the TRP domain were experimentally characterized in this study, or in reference (36). The location of the W682 residue in TRPM8 that we experimentally characterized is also noted here in cyan. The dual numbering S4-S5 loop and in the TRP domain indicates the difference in numbering between the rTRPM8 that we use in experiments and the fcTRPM8 used in cryoEM studies. The MHR4 region in TRPM8 that contains the K605 PI(4,5)P_2_ contact residue in TRPM8 is not shown, as its sequence is not conserved in TRPM3. (B-M) Human TRPM3_1325_ splice variant, or its mutants were expressed in *Xenopus* oocytes, and two electrode voltage clamp (TEVC) experiments were performed to measure the activity of TRPM3 as described in the Methods section using a ramp protocol from –100 mv to +100 mv. Currents induced by 50 μM PregS were measured before and after treatment with 35 μM wortmannin to deplete PI(4,5)P_2_. (B,C) Representative traces of wild type TRPM3 (B), and K992A mutant (C) before (*left*) and after (*right*) treatment of the same oocyte with 35 μM wortmannin for 2 hours. Top traces show currents at +100 mV; dash lines indicate 0 current; bottom traces show currents at −100 mV. Applications of 50 μM PregS are indicated by red lines. (D-H) Data summary of the percentage inhibition of PregS-induced currents by wortmannin treatment at 100 mV for different mutants: W761F (D), N991A (E), K992A (F), R1131Q and R1141Q (G) and K1128Q (H). (I-M) Current amplitudes of various mutants at 100 mV. Each symbol represents measurement of one oocyte from 2 to 3 independent preparations. Statistical significance was calculated with t-test, or one way ANOVA (G,L), p values are shown on bar graphs.

The PI(4,5)P_2_ binding site in TRPM3 is formed by parts of the preS1 segment, the S4-S5 loop and the proximal C-terminal TRP domain of the same subunit. The closest contact residues with PI(4,5)P_2_ are W761 in the preS1 segment, the N991and K992 residues in segment connecting the voltage-sensor like domain (VSLD, S1-S4) to the S4-S5 linker and the R1131 in the TRP domain (**Fig. 1E**). **Figure 1E** also shows the location of two additional residues, K1128 and R1141 in the TRP domain, which are not in close contact with PI(4,5)P_2_, but we experimentally characterized their mutations, see later. The numbering of these residues corresponds to the splice variant of hTRPM3 (hTRPM_1325_) (15,38), which we used in the majority of our experiments.

Comparing our model of TRPM8 in complex with PI(4,5)P_2_ with the subsequently determined two cryo-EM structures of TRPM8 with PI(4,5)P_2_ (28) with icilin (6NR3), and with the menthol analog WS12 (6NR2), offered *a posteriori* validation of our modeling (**Table S1**). **Figure S3** compares the PI(4,5)P_2_ binding pockets of the TRPM8-PI(4,5)P_2_-icilin structure, the TRPM8-PI(4,5)P_2_-WS12 structure, and our computational model. Our model superimposes very well with both structures in the transmembrane (TM) domain, but it shows a better alignment of the PI(4,5)P_2_ binding site with the TRPM8-PI(4,5)P_2_-WS12 (6NR2) structure. Interestingly, the structural difference between the two experimental structures is larger than that observed between our model and the experimental complexes. Yin et al. listed five key residues in their cryo-EM co-structures critical for PI(4,5)P_2_ interaction: R997 in the TRP domain, R850 in the S4-S5 loop, N692 and R688 in the pre-S1 segment (**Fig. 3A**) and K605 in the neighboring N-terminal cytoplasmic MHR4 domain (28). All these residues, with the exception of R850, are in contact with, or very close to PI(4,5)P_2_ in our TRPM8-PI(4,5)P_2_ model. R850 is in contact with the acyl chain of PI(4,5)P_2_ in our model, and only in contact with the PI(4,5)P_2_ headgroup in the 6NR3, but not in the 6NR2 structure, which is consistent with the better alignment of our model with the 6NR2 PI(4,5)P_2_ TRPM8 structure. Overall, our TRPM8-PI(4,5)P_2_ docking model validates our computational approach to identify the TRPM3 PI(4,5)P_2_ binding site and suggest that PI(4,5)P_2_ likely binds to a site that is similar in TRPM3 and TRPM8.

Furthermore, superimposition of our model to the experimental structure of TRPM7 in EDTA (5ZX5) (33) reveals that the docked PI(4,5)P_2_ in our model of TRPM3 fits well in a cavity of the experimental structure of TRPM7 that accommodates a detergent cholesteryl hemisuccinate (CHS) molecule (**Fig. S4**). Whether this binding site is occupied by PI(4,5)P_2_ in TRPM7 in a cellular environment, remains to be determined, nevertheless the presence of this lipid binding pocket in TRPM7 suggests that the location of the PI(4,5)P_2_ binding site may be conserved in multiple members of the TRPM sub-family.

In our TRPM3-PI(4,5)P_2_ model, residues K992 and R1131 are equivalent to the experimentally determined PI(4,5)P_2_ contact sites R850 and R997 in TRPM8, located in the S4-S5 loop and the TRP domain (**Fig. 3A**). Specifically, N991 is adjacent to K992, and W761 in the pre-S1 segment of TRPM3 is shifted 6 residues from the R688 residue in TRPM8. The equivalent of W761 in TRPM8 (W682) is relatively close to PI(4,5)P_2_ in TRPM8, and so is the equivalent of R688 in TRPM3 (M767) highlighting the generally similar importance of the pre-S1 segment in PI(4,5)P_2_ binding in the two channels. The largest difference between the two binding sites is that the MHR4 region, which carries K605 in TRPM8, is not conserved in TRPM3, and the equivalent residue is far away from PI(4,5)P_2_ in our TRPM3 model. Overall, the two channels bind PI(4,5)P_2_ in a similar, yet not identical manner (**Fig. S5**).

Next, we mutated the predicted PI(4,5)P_2_ interacting residues in TRPM3 and tested the effects of the mutations on sensitivity to inhibition by decreasing PI(4,5)P_2_ levels. We expressed the wild type and mutant channels in Xenopus oocytes, and performed two electrode voltage clamp experiments. We stimulated channel activity with 50 μM pregnenolone sulfate (PregS) and measured current amplitudes, and then incubated the oocytes with 35 μM wortmannin for 2 hours, and measured PregS-induced currents in the same oocytes (**Fig. 3B,C**). Wortmannin at this concentration inhibits phosphatidylinositol 4-kinases (PI4K), and has been used to inhibit the activity of PI(4,5)P_2_ dependent ion channels (39). Mutating a PI(4,5)P_2_ interacting residue is expected to increase inhibition by wortmannin (39). We mutated the TRP domain positively charged residues to Q, as equivalent mutations in TRPM8 were shown to be functional, and affect PI(4,5)P_2_ interactions (36). The rest of the resides we mutated to A, but the W761A mutant was non-functional, thus we functionally characterized W761F instead. Mutations of all computationally predicted PI(4,5)P_2_ interacting residues (W761F, N991A, K992A, and R1131Q) showed significantly higher inhibition after wortmannin treatment than wild type TRPM3 (**Fig. 3D-G**), and their current amplitudes were also significantly lower than wild type TRPM3 (**Fig 3I-L**). We also generated two additional mutations in the TRP domain in residues that are not in contact with PI(4,5)P_2_, K1128Q and R1141Q. Both mutants showed similar current amplitudes to wild type TRPM3 (**Fig. 3L,M**). The K1128Q mutant showed similar inhibition to WT (**Fig. 3H**), but the R1141Q mutant showed a small but significant increase in wortmannin inhibition compared to wild type (**Fig. 3G**). This mutation is equivalent to R1008Q in the rat TRPM8, which reduced both PI(4,5)P_2_ and menthol sensitivity (36), but it was in contact with the menthol analogue WS12, but not with PI(4,5)P_2_ in the cryoEM structure of fcTRPM8 (R1007) (28). Therefore, it is possible that this mutation affected PI(4,5)P_2_ sensitivity indirectly.

Next we confirmed our data in whole cell patch clamp experiments using the rapamycin-inducible 4’ 5’ phosphoinositide phosphatase pseudojanin (40). Figure S6A-C shows that 100 nM rapamycin induced a significantly higher inhibition of the N993A mutant of the mouse TRPM3α2 (equivalent of N991A in hTRPM3) than the wild type mTRPM3α2 when the channels were stimulated with 25 μM PregS. Next, we stimulated the mTRPM3α2 with the combination of 25 μM PregS and 10 μM clotrimazole, which was shown to open an alternative pore, characterized by less prominent outward rectification (41). Application of 100 nM rapamycin induced a significantly larger inhibition of currents induced by clotrimazole plus PregS in the N993A mutant compared to the wild type mTRPM3α2 (**Fig S6D-F**).

Mutation of the PI(4,5)P_2_ contact site R998Q resulted in a right shift in the diC_8_ PI(4,5)P_2_ dose response in excised patches (36). TRPM3 currents in excised patches show a less steep concentration dependence, with no clear saturation at higher concentrations (21). This and the low current amplitudes in the mutants prevented us from reliably comparing PI(4,5)P_2_ does responses in our mutants. It was reported for TRPV1 that mutating a PI(4,5)P_2_ interacting residue increased the relative efficiency of PI(4)P to stimulate channel activity compared to PI(4,5)P_2_ (42). Therefore, we tested the relative effects of PI(4)P and PI(4,5)P_2_ on the N991A mutant. **Figure S7** shows that the relative effect of PI(4)P compared to PI(4,5)P_2_ did not change.

Next, we used MM/GBSA calculations to predict the changes in the binding free energy (ΔΔG) of PI(4,5)P_2_ to the native (wild type) TRPM3 versus the mutant channels that were characterized in **Figure 3**. As shown in **Figure 4,** the binding of PI(4,5)P_2_ is guided by a number of stabilizing interactions established with key contact residues (**Fig. 4B and Table S2**). Mutations of all these residues in our model resulted in a decreased PI(4,5)P_2_ binding affinity (for a native protein to bind better than the mutant, the calculated ΔΔG value is positive). Specifically, K992A had a more prominent effect than W761F, N9991A or R1131Q. This correlates well with K992A also having the most pronounced effect on inhibition by PI(4,5)P_2_ depletion (**Fig. 3F**). Regarding the binding modes, K992 engages in multiple interactions with PI(4,5)P_2_, including two hydrogen bonds and two salt bridges. Mutating K992 to alanine resulted in the loss of all these interactions, with the exception of one hydrogen bond (**Fig. 4 G-H**). Similar behavior was observed with the mutation R1131Q, the contact residue exerting the second largest effect on binding affinity, which resulted in the loss of one hydrogen bond and two salt bridges (**Fig. 4 I-J**). Next, mutating W761 to phenylalanine and N991 to alanine resulted in the loss of one hydrogen bond interaction each (**Fig. 4 C-D** and **Fig. 4 E-F,** respectively). The R1128Q mutation had only a very small effect on both ΔΔG and the related binding mode, which correlates well with it not being in close contact with PI(4,5)P_2_ in our model, and the lack of effect on wortmannin inhibition. The R1141Q mutant, which is also not a PI(4,5)P_2_ contact site, also had only a minimal effect on both ΔΔG and the related binding mode, indicating that the small, but significant effect on wortmannin inhibition was likely due to indirect effects. Overall, all Δ affinity calculations supported our computational docking and agreed with the experimental functional characterization of the PI(4,5)P_2_ interacting residues.

**Figure 4.**
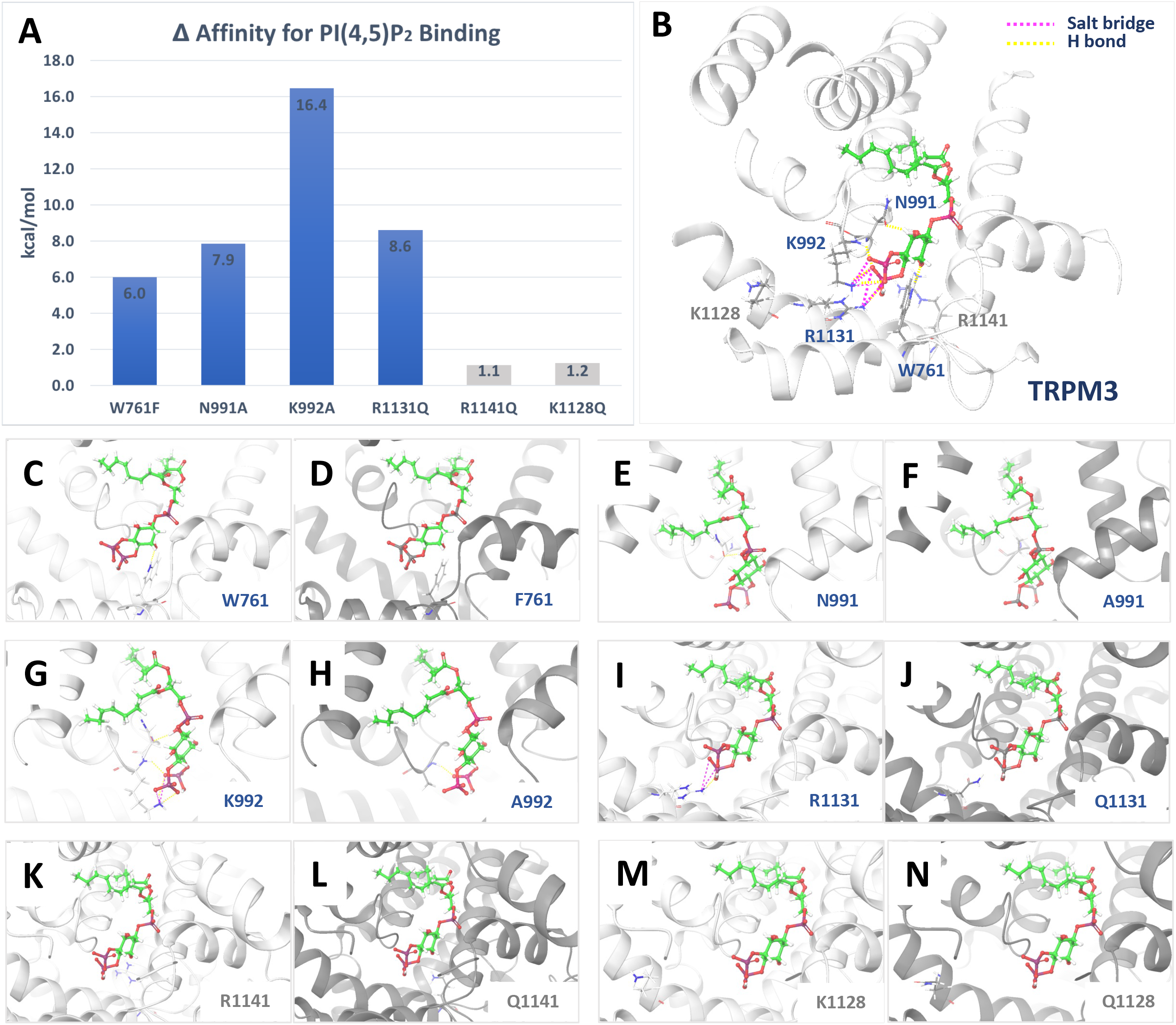
Effect of mutations in the PI(4,5)P_2_ binding site residues of TRPM3. MM/GBSA calculations were performed as described in the Methods section. (**A**) Change in binding affinity (ΔΔG; kcal/mol) upon mutating protein residues *in silico* in the PI(4,5)P_2_ binding site of TRPM3. In blue, mutants that bind significantly worse than the native protein, indicating loss of interaction with PI(4,5)P_2_ upon mutations. In grey, mutants with no significant effect on binding. (**B to N**) Binding of PI(4,5)P_2_ to native and mutant TRPM3 channels. In (B), native TRPM3. In (C to N), the effect of each individual mutation is analyzed. Protein atoms are shown in new cartoon representation, in white and grey color for native and mutant channels, respectively. Ligand atoms are in ball- and-stick representation, with C, O, H and P atoms colored in green, red, white and yellow, respectively. Hydrogen bonds and salt bridges are represented as yellow and pink dotted lines, respectively.

Next, we mutated two residues in the rTRPM8 that are equivalent to PI(4,5)P_2_ interacting residues in our TRPM3 model. The R851 residue in TRPM8 corresponds to the K992 residue in the S4-S5 linker in TRPM3 (**Fig. 3A**), and it was in direct contact with PI(4,5)P_2_ in the cryo-EM structure of the fcTRPM8 (R850) (28). The W682 residue is the equivalent of W761 in TRPM3 (**Fig. 3A**), and while is not in a direct contact with PI(4,5)P_2_ in the fcTRPM8 cryo-EM structure (R850) it is located relatively close. Since the W682A mutant was non-functional, we characterized the W682Q, which displayed small, yet measurable menthol-induced currents. **Figure 5A-G** shows that both the R851Q and the W682Q mutants showed significantly higher level of inhibition by wortmannin, with W682Q having a larger effect. Current amplitudes showed a similar pattern, both mutants were significantly lower that wild type TRPM8, and the W682Q having a larger effect (**Fig. 5H**). Wortmannin treatment substantially accelerated deactivation after cessation of menthol stimulation (**Fig. 5A-F**), which is in contrast to TRPM3, where the deactivation kinetics after washing out PregS was not affected by wortmannin (**Fig. 3B,C**).

**Figure 5.**
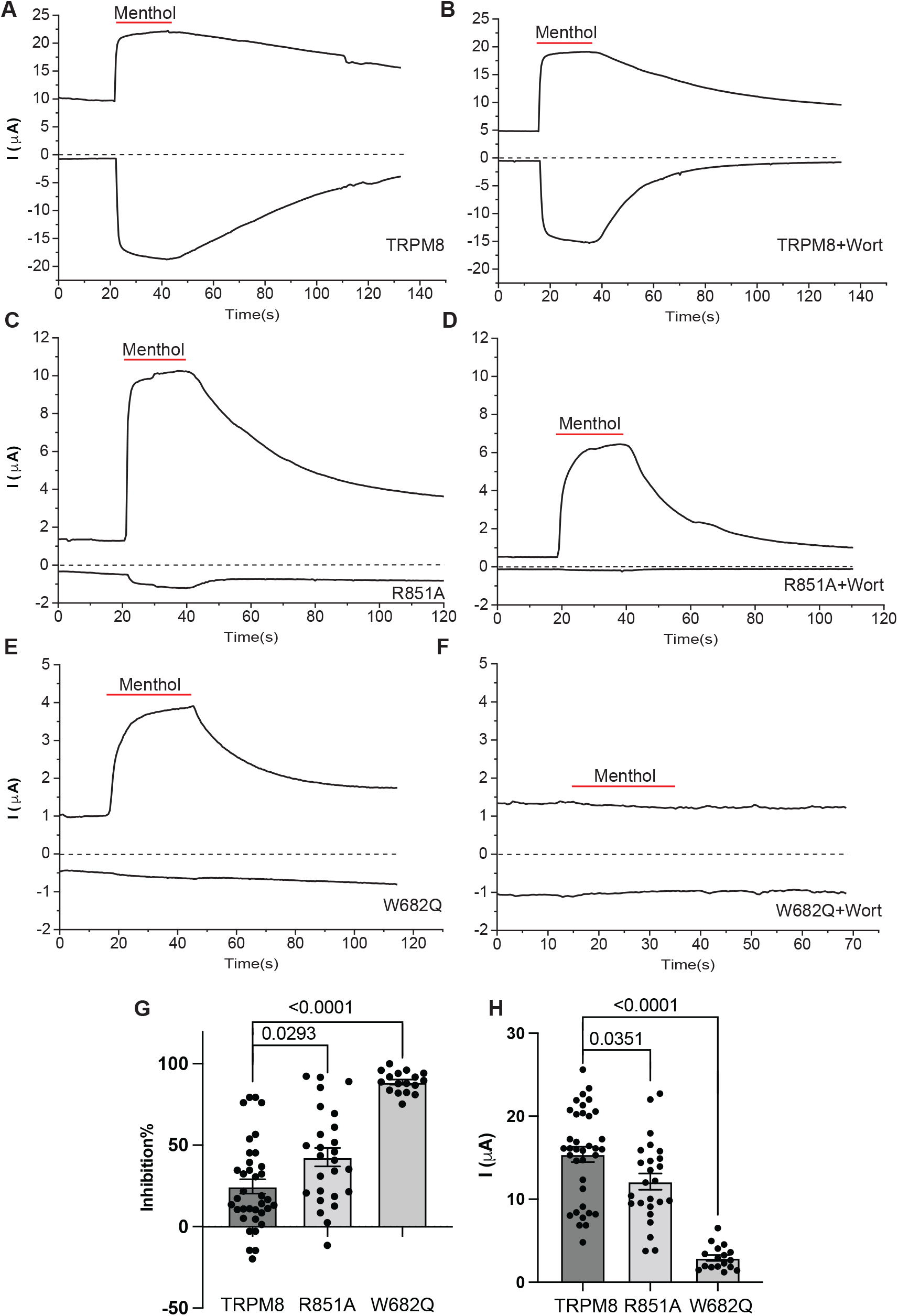
Mutating putative PI(4,5)P_2_ interacting residues increases sensitivity of TRPM8 to inhibition by PI(4,5)P_2_ depletion. Rat TRPM8 or its mutants were expressed in Xenopus oocytes. TEVC experiments were performed as described in the Methods section using ramp protocol from –100 mv to +100 mv. Menthol (500 μM) was applied to activate TRPM8 channels and 35 μM wortmannin was applied for 2 hours to deplete PI(4,5)P_2_. (A-F) Representative traces of TRPM8 before (A) and after wortmannin treatment (B), R851A before (C) and after wortmannin treatment (D) and W682Q before (E) and after wortmannin treatment (F). Top traces show currents at 100 mV; dash lines indicate 0 current; bottom traces show currents at −100 mV. Applications of 500 μM menthol are indicated by red lines. (G) Percentage inhibition induced by PI(4,5)P_2_ depletion in the TRPM8 wild type group at −100 and 100 mV. (H) Percentages of inhibition evoked by PI(4,5)P_2_ depletion at 100 mV plotted for wild type, R851A and W682Q. (I) Current amplitudes of all three groups at 100 mV. Each symbol represents an individual oocyte. All experiments were from 2 to 3 independent preparations. P values are shown on bar graphs. Statistical significance was calculated with t-test (G), or one way ANOVA (H,I). P values are shown on bar graphs.

Stimulation with menthol, or cold, was shown to increase the apparent affinity of TRPM8 for PI(4,5)P_2_ (36) indicating an allosteric interaction between menthol and PI(4,5)P_2_ activation. Next, we asked if this allosteric interaction also happens in the opposite direction, and tested if mutations of the PI(4,5)P_2_ interacting residues in TRPM8 has an effect on agonist sensitivity. **Figure 6** shows that both the R851A and the W682Q mutant had right shifted menthol dose response. Similar to the effect on current amplitudes and wortmannin inhibition, the effect of the W862 mutant was more pronounced than that of the R852A. We also tested if a similar allosteric effect also exists in TRPM3. **Figure 7** shows that both the N991A and the K992A mutant shifted the PregS dose response to the right.

**Figure 6.**
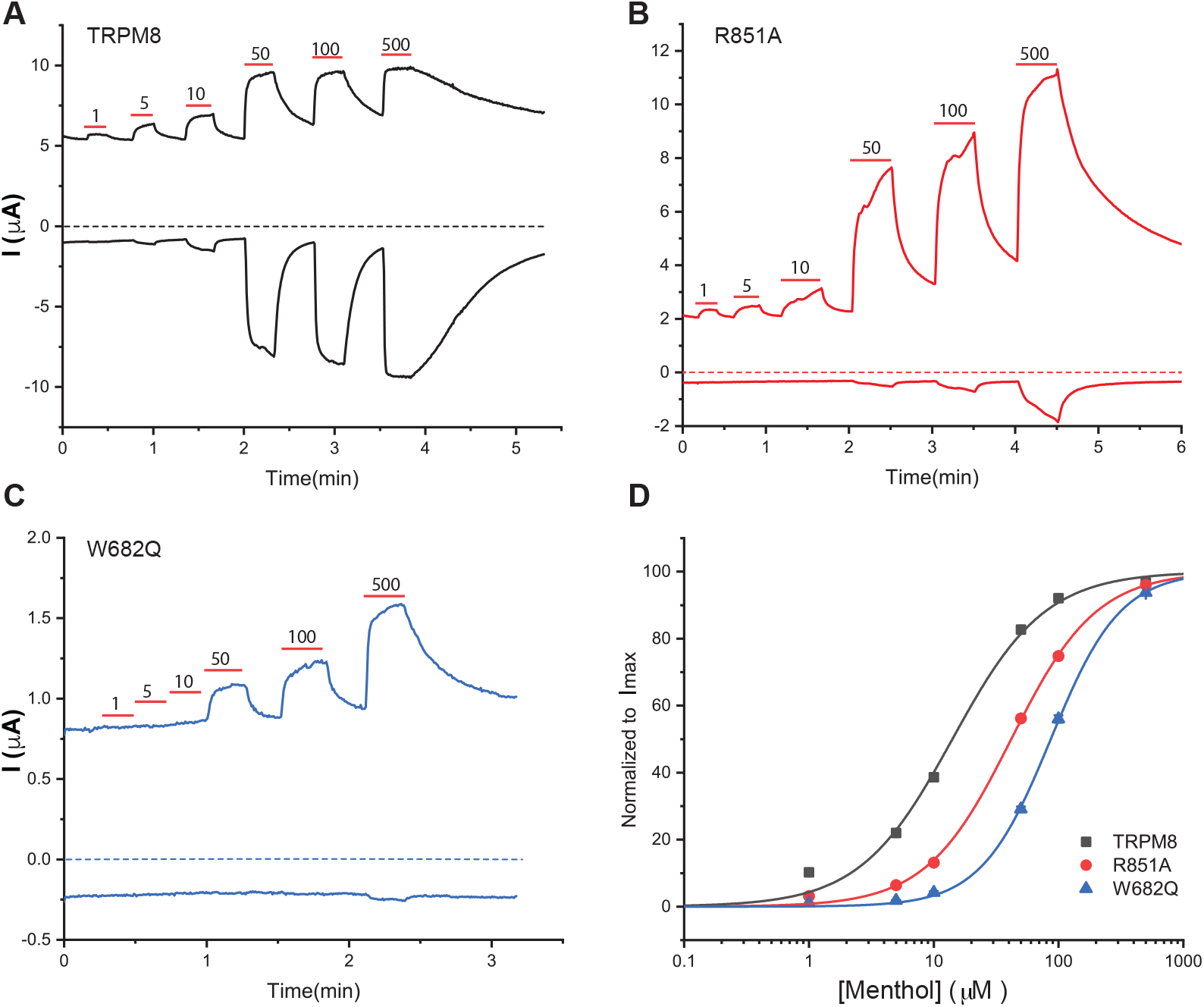
Mutating putative PI(4,5)P_2_ interacting residues decreases sensitivity of TRPM8 to menthol activation. TEVC experiments were performed using a ramp protocol from −100 mV to 100 mV, as described in the Methods section. (A-C) Representative traces of TRPM8 (A), R851A (B) and W682Q (C). Top traces show currents at +100 mV; dash lines indicate 0 current; bottom traces show currents at −100 mV. Applications of various concentrations of menthol (μM) are indicated by red lines. (D) Hill1 fit of the concentration dependence of menthol at 100 mV for TRPM8 and mutated channels. Symbols represent mean ± SEM for n=12-13 measurements from 2 different oocyte preparations.

**Figure 7.**
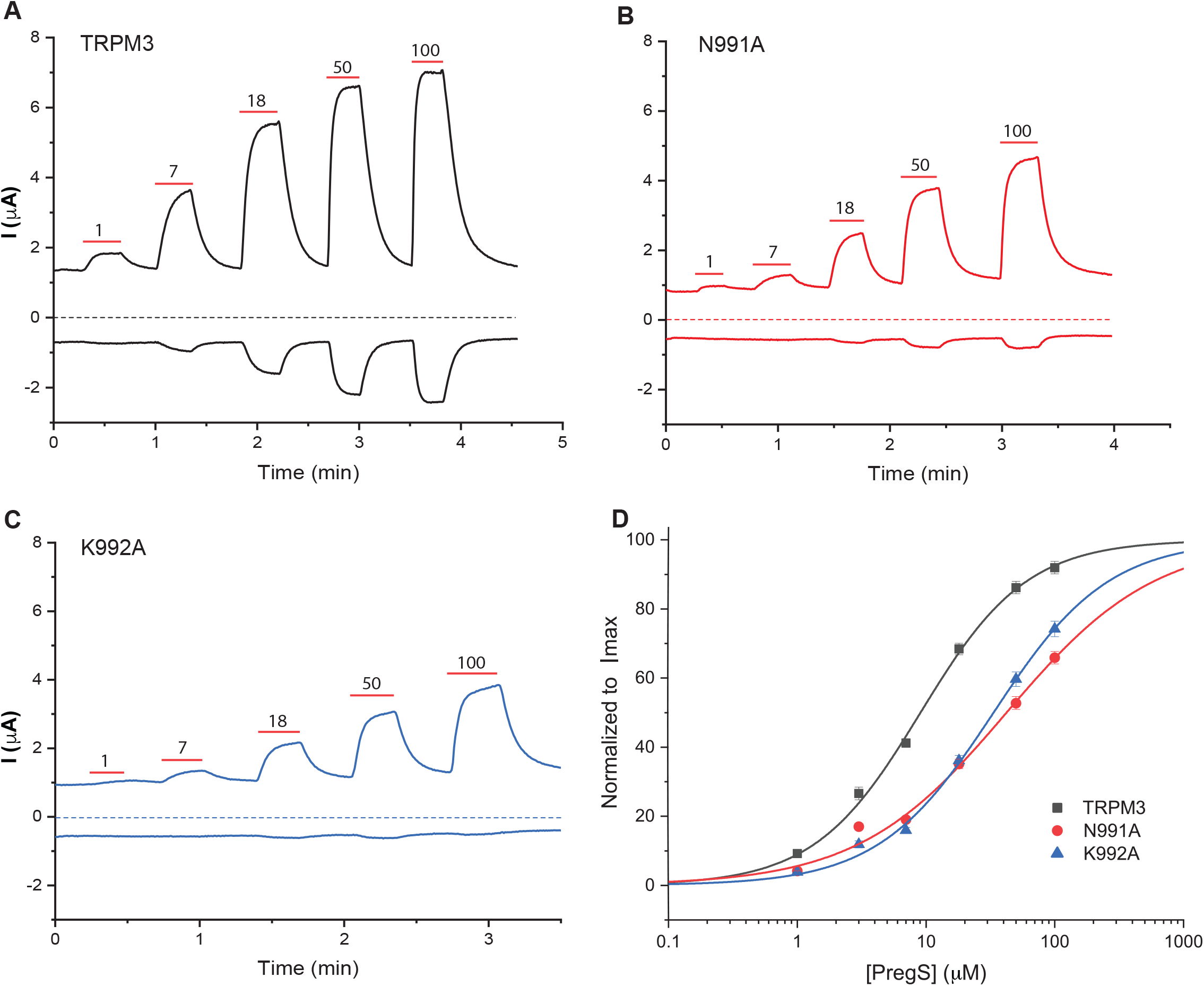
Mutating putative PI(4,5)P_2_ interacting residues decreases sensitivity of TRPM3 to PregS activation. cRNA of either the TRPM3_1325_ splice variant, or its mutants were injected into Xenopus oocytes. TEVC was performed to measure TRPM3 currents using a ramp protocol from − 100 mV to 100 mV, as described in the Methods section. (A-C) Representative traces of hTRPM3 wild type (A), N991A (B) and K992A (C). Top traces show currents at +100 mV; dash lines indicate 0 current; bottom traces show currents at −100 mV. Applications of various concentrations of PregS (μM) are indicated by red lines. (D) Hill1 fit of the concentration dependence of PregS at 100 mV for TRPM3 and mutated channels. Symbols represent mean ± SEM for n=12-13 measurements from 2 different oocyte preparations.

TRPM3 activity is inhibited by direct binding of Gβγ to the channel (9). To test if an allosteric interaction between Gβγ inhibition and PI(4,5)P_2_ activation is present, we expressed wild type and mutant TRPM3 channels with, or without Gβ1γ2 in Xenopus oocytes, and measured PregS induced currents. The N991A and K992A mutants were inhibited significantly more by Gβγ than wild type TRPM3 (**Fig. 8A-E**). The K1128Q mutant, which did not affect PI(4,5)P_2_ sensitivity, had no effect on Gβγ inhibition either (**Fig 8E**). Interestingly, the R1131Q mutant was not inhibited, rather potentiated by coexpressing Gβγ (**Fig. 8E**). Consistently with the lack of inhibition by Gβγ, the R1131Q mutant was also not inhibited by stimulating Gi-coupled M2 muscarinic acetylcholine receptors (**Fig 8F-J**). These data indicate that while there is an allosteric interaction between PI(4,5)P_2_ and Gβγ, the R1131 residue in the TRP domain also play some role in transmitting the inhibitory effect of Gβγ.

**Figure 8.**
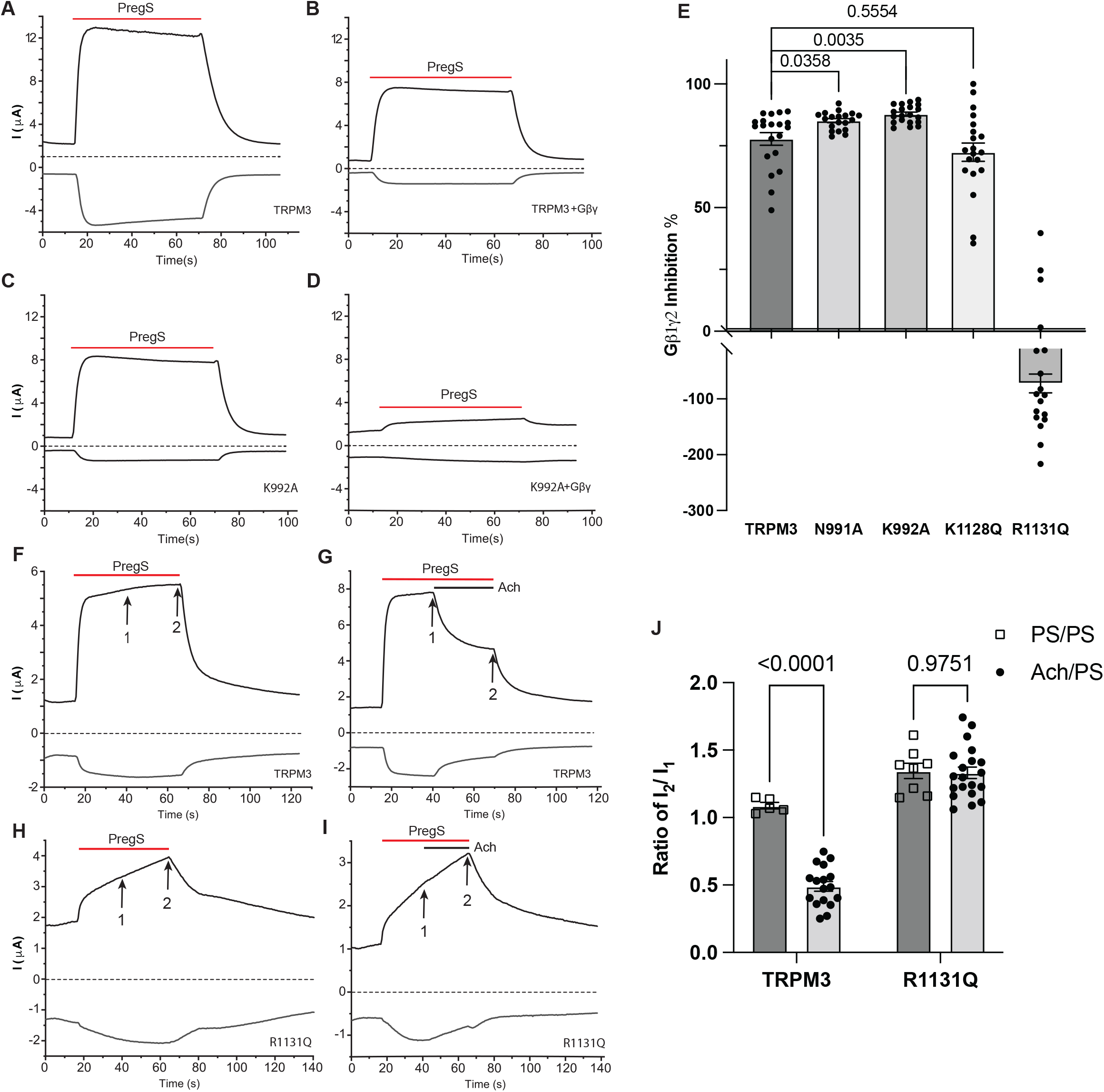
Relationship between Gβγ and PI(4,5)P_2_ regulation of TRPM3. hTRPM3 or its mutants were expressed in oocytes with or without Gβ1γ2 subunits for panels A-E, or with muscarinic hM2 receptors for panels F-J. TEVC was used to measure channel activity using a ramp protocol from −100 mV to 100 mV as described the Methods section. (A-D) Representative traces of TRPM3 (A), TRPM3 co expressed with Gβγ subunits (B), K992A (C) and K992A co expressed with Gβγ subunits (D). (E) Data summary shows the inhibition caused by Gβγ subunits in different mutant groups. Percentages of inhibition was calculated by normalizing decreased current amplitudes to the average currents induced by 50 μM PS in control oocytes without Gβγ subunits. (F-J) Representative traces of TRPM3 treated with PregS alone (F), TRPM3 treated with PregS and acetylcholine (G), R1131Q treated with PregS alone (H) and TRPM3 treated with PregS and acetylcholine (I). Top traces show currents at +100 mV; dash lines indicate 0 current; bottom traces show currents at −100 mV. Application of 50 μM PregS is indicated by red lines and application of 5 μM acetylcholine is indicated by black lines. The 1^st^ arrow indicates currents induced by PregS and the 2^nd^ arrow indicates currents after application of acetylcholine or vehicle with continuous application of PregS. (J) Data summary shows the ratio of current amplitudes between points 2 and 1 indicated on the representative traces. Each symbol represents individual oocyte from three (E) and two (J) independent preparations. Statistical significance was calculated with one way ANOVA (E), and two-way ANOVA (J). P values are shown on bar graphs.

Gain of function mutations in TRPM3 have recently been shown to cause intellectual disability and seizures (13–15). The two disease-associated mutations, V990M and P1090Q were shown to increase basal channel activity, as well as increase agonist sensitivity and increased heat sensitivity, with V990Q affecting agonist sensitivity more prominently, while P1090Q predominantly affecting heat sensitivity (15). Next, we tested if the increased basal activity and agonist sensitivity also translates into higher sensitivity to PI(4,5)P_2_. **Figure 9** shows that when wild type and V990Q and P1090Q mutant channels were treated with 35 μM wortmannin for 2 hours, the three groups were inhibited to a similar extent. They were also inhibited to a similar extent when currents were evoked by the respective EC_50_ of the mutant and wild type channels. These data indicate that the disease mutants do not increase channel activity by increasing their apparent affinity for PI(4,5)P_2_.

**Figure 9.**
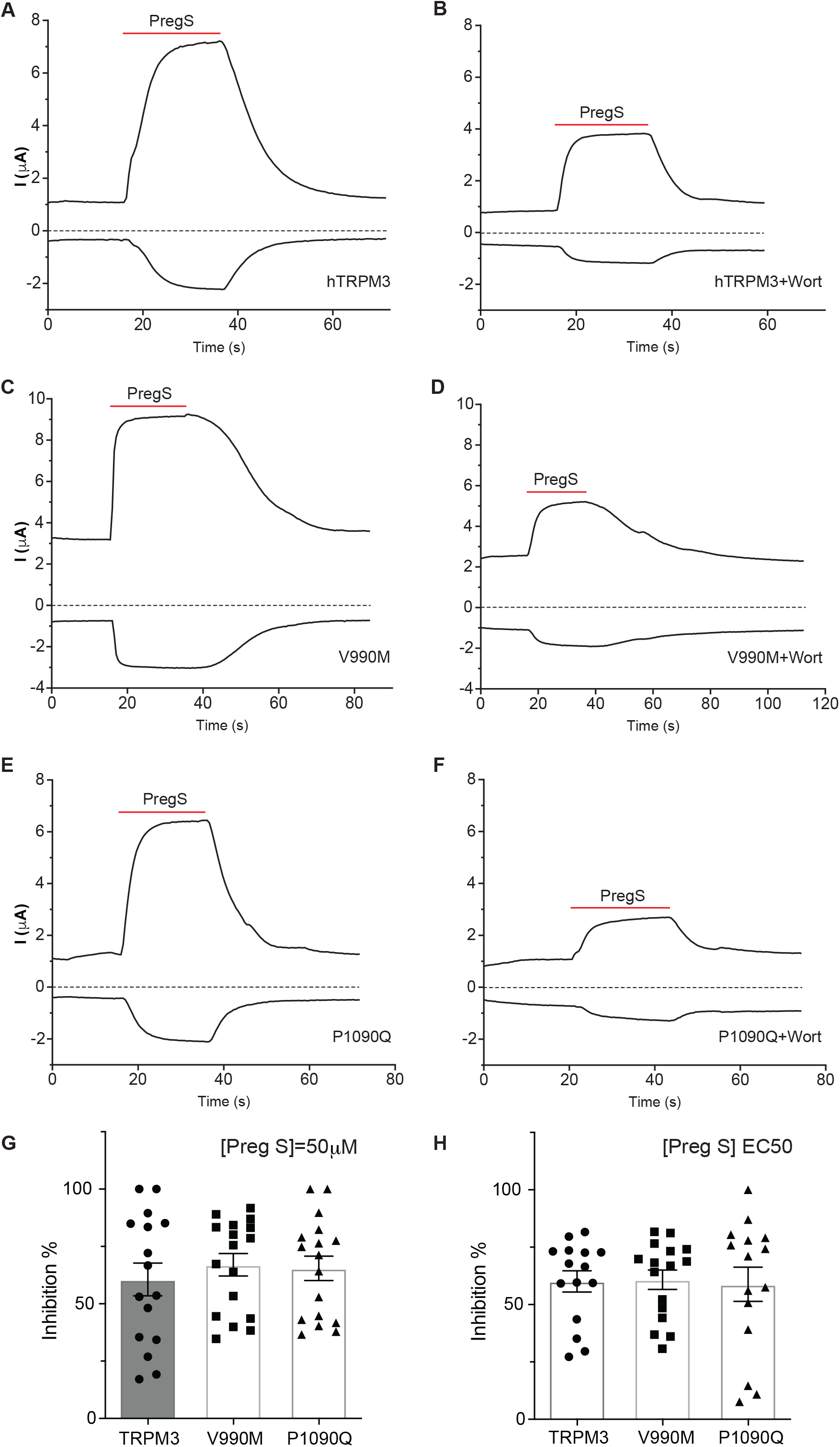
Disease associated gain of function mutants do not change the TRPM3 sensitivity to the PI(4,5)P_2_ depletion. hTRPM3, V990M or P1090Q were expressed in oocytes and 35 μM wortmannin (2h) was used to deplete PI(4,5)P_2_. TEVC was performed as described in the Methods section. (A-F) Representative traces of hTRPM3 (A), hTRPM3 after wortmannin treatment (B), V990M (C), V990M after wortmannin treatment (D), P1090Q (E) and P1090Q after wortmannin treatment (F). Top traces show currents at +100 mV; dash lines indicate zero current; bottom traces show currents at −100 mV. Applications of 50 μM PregS is indicated by red lines. (G) Data summary of wortmannin inhibition of currents induced by 50 μM PregS. (H) Data summary of wortmannin inhibition of currents induced by EC_50_ concentrations of PregS, 17 μM, 0.6 μM and 7 μM for hTRPM3, V990M and P1090Q respectively. Symbols represent individual oocytes from three (G) and two (H) different preparations.

## DISCUSSION

Our work aims to understand the molecular mechanism of PI(4,5)P_2_ regulation of TRPM3. We used computational docking, changes in the binding affinity estimated by computational mutagenesis, site directed mutagenesis and electrophysiology to identify PI(4,5)P_2_ interacting residues in the channel protein. Our data indicate that residues in three regions, the pre-S1 segment, the S4-S5 loop, and the TRP domain play important roles in forming the PI(4,5)P_2_ binding site in TRPM3. Mutation of PI(4,5)P_2_ interacting residues decreased the binding affinity *in silico* (positive ΔΔG values in **Fig. 4** indicate that the native protein binds better than the mutant), and increased sensitivity to inhibition by decreasing PI(4,5)P_2_ levels in electrophysiology experiments (**Fig. 3**). Mutating PI(4,5)P_2_ interacting residues also decreased sensitivity to PregS activation, and increased sensitivity to Gβγ inhibition indicating allosteric interaction between PI(4,5)P_2_ and agonists as well as a physiological inhibitor. On the other hand, disease-associated gain of function mutations did not change PI(4,5)P_2_ sensitivity, indicating that the mutations did not increase channel activity by enhancing PI(4,5)P_2_ activation.

There are currently five channels in the TRPM family for which structural data are available (43): TRPM2 (44), TRPM4 (45), TRPM5 (46), TRPM7 (33) and TRPM8 (28). While all of these channels have been shown to be positively regulated by PI(4,5)P_2_ (23), only TRPM8 has a co-structure with this lipid (28). When we compare the experimentally determined PI(4,5)P_2_ binding site in TRPM8 with our computationally identified binding site in TRPM3, the two overlap, sharing some of the interacting residues, with some differences (**Fig 3A, Fig. S5**). Overall, the preS1 segment, the S4-S5 loop and the TRP domain are involved in both channels in forming the PI(4,5)P_2_ binding site. The R1131 residue in the TRP domain in TRPM3 is equivalent to the R997 PI(4,5)P_2_ contact residue in the fcTRPM8 structure (28), and to the R998 residue in the rat TRPM8, which was proposed as a PI(4,5)P_2_ interacting residue and experimentally shown to exhibit decreased PI(4,5)P_2_ sensitivity before structures became available (36). The K992 residue in the S4-S5 loop of TRPM3 is equivalent to the R850 PI(4,5)P_2_ contact residue in the fcTRPM8 structure, and to the R851 residue in the rat TRPM8 that we characterized in this study (**Fig. 5, Fig. 6**). The preS1 segment of the fcTRPM8 has two PI(4,5)P_2_ contact residues R688 and N692 (**Fig. 3A**). These residues are not conserved in TRPM3 (**Fig. 3A**); yet the equivalent residues in our TRPM3 model – i.e., M767 and G672 – are both located within 5 Å of the PI(4,5)P_2_ head-group (**Fig. S5D**), with M767 engaging hydrophobic interactions that stabilize the overall complex. The W761 PI(4,5)P_2_ contact residue in the pre-S1 of TRPM3, is equivalent to the W682 residue in TRPM8, which is close to the R688 residue as well as to PI(4,5)P_2_, but it was not close enough to designate it as a PI(4,5)P_2_ contact site in TRPM8 (28). Interestingly, when we mutated this residue to a glutamine (W682Q) in the rat TRPM8, it behaved similar to the W761F mutation in TRPM3, i.e. it increased sensitivity to PI(4,5)P_2_ depletion (**Fig. 5**). Whether this residue is in a closer contact with PI(4,5)P_2_ in a cellular environment in the rat TRPM8, or its mutation affected PI(4,5)P_2_ interactions indirectly, or both, it is difficult to tell. Finally, the K605 residue from an adjacent cytoplasmic MHR4 domain was also in contact with PI(4,5)P_2_ in TRPM8. This residue is not conserved in TRPM3, and was not close to PI(4,5)P_2_ in our homology model.

It is well established that channel agonists can increase PI(4,5)P_2_ sensitivity (apparent affinity) for various PI(4,5)P_2_ sensitive ion channels. For example, the apparent affinity of the G-protein activated inwardly rectifying K^+^ channel GIRK4 (Kir3.4) for PI(4,5)P_2_ is increased by factors that stimulate channel activity, such as Gβγ and intracellular Na^+^ (47). The apparent affinity of TRPM8 for PI(4,5)P_2_ was shown to be increased by both cold, and menthol (36), and the apparent affinity of TRPV1 for PI(4,5)P_2_ activation was increased by capsaicin (48). The opposite was also proposed, as a mutation in the putative PI(4,5)P_2_ interacting residue R1008 in TRPM8 not only decreased apparent affinity for PI(4,5)P_2_, but also induced a marked right shift in the menthol dose-response (36). In the view of the structures of TRPM8 however, this residue is likely to be a menthol interacting residue, as it was in close contact in the TRPM8 structure with the menthol analogue WS12, but not with PI(4,5)P_2_ (28), therefore it most likely primarily affected menthol sensitivity, and the effect on PI(4,5)P_2_ was a secondary allosteric effect. Our data indicate that in both TRPM8 and in TRPM3, mutating PI(4,5)P_2_ contact residues also decreases agonist sensitivity. Similarly, mutating most PI(4,5)P_2_ interacting residues also made it easier for TRPM3 to be inhibited by Gβγ. This is likely to be an allosteric effect, as the Gβγ binding peptide in TRPM3 (12) is located on the outer surface of the channel, far away from the PI(4,5)P_2_ binding site (**Fig. S8**). Interestingly, the R1131Q mutant did not display any Gβγ inhibition, pointing to the complex role of this residue in channel regulation.

In contrast to the apparent allosteric interaction between PI(4,5)P_2_ and agonist or Gβγ, disease-associated gain of function mutations in TRPM3 that increased both heat and agonist sensitivity (15), did not decrease sensitivity for inhibition by PI(4,5)P_2_ depletion (**Fig. 9**.), indicating that the mechanism of increased channel activity is not the consequence of increased sensitivity to PI(4,5)P_2_.

In an earlier work, well before structures became available, three residues in the TRP domain of TRPM8 were proposed to act as PI(4,5)P_2_ interacting residues (36). While mutations in all three of them decreased PI(4,5)P_2_ apparent affinity (36), only one of them was in direct contact with PI(4,5)P_2_ in the TRPM8 PI(4,5)P_2_ structures that were determined later (28). Before TRP channel structures became available, a short “PH-domain-like” segment with several positively charged residues was proposed to act as a PI(4,5)P_2_ interaction site in TRPM4, (49). Even though mutations in this segment behaved in a way compatible with reduced PI(4,5)P_2_ interactions, this segment was far away from the plasma membrane in the subsequently determined TRPM4 structures, which is incompatible with acting as a PI(4,5)P_2_ interacting domain (45). These findings indicate that in the absence of structural information it is difficult to predict PI(4,5)P_2_ interacting residues with high accuracy.

In our earlier work, we used a homology model, based on the structure of TRPV1 combined with mutagenesis, to predict PI(4,5)P_2_ interacting residues in the epithelial Ca^2+^ channel TRPV6 (50). Our homology model-based PI(4,5)P_2_ site was very similar to the experimentally determined PI(4,5)P_2_ binding site in TRPV5, with the same key contact residues (50,51). TRPV5 and TRPV6 are products of a relatively recent gene duplication, and they share 75% identity, and they are functionally far more similar to each other than to other members of the TRPV subfamily. This gives us confidence that our computationally determined PI(4,5)P_2_ binding site in TRPV6 likely reflects the actual PI(4,5)P_2_ binding site with a reasonable accuracy, even if there is no TRPV6 PI(4,5)P_2_ co-structure available currently. Similarly, in the current work, we docked PI(4,5)P_2_ to the apo structure of TRPM8 (30), which showed a high level of overlap with the PI(4,5)P_2_ binding site of TRPM8 determined subsequently by cryoEM studies (28). This makes us confident that our experimentally tested computational prediction of the PI(4,5)P_2_ binding site in TRPM3 reflects the functionally relevant PI(4,5)P_2_ binding site with reasonable accuracy.

In conclusion, our data provides mechanistic insight into regulation of TRPM3 by its key physiological co-factor, PI(4,5)P_2_. We identify its binding site on the channel, characterize the interaction between PI(4,5)P_2_ and other physiological regulators of TRPM3, and compare its regulation by PI(4,5)P_2_ to that of TRPM8.

## METHODS

### *Xenopus laevis* oocytes preparation and RNA injection

All procedures of preparing *Xenopus laevis* oocytes were approved by the Institutional Animal Care and Use Committee at Rutgers New Jersey Medical School. Frogs were anesthetized in 0.25% ethyl 3-aminobenzoate methanesulfonate solution pH 7.4 (MS222; Sigma-Aldrich), then bags of ovaries were surgically collected from and rotated with 0.1-0.3 mg/ml type 1A collagenase (Sigma-Aldrich) at 16 °C overnight in OR2 buffer containing 82.5 mM NaCl, 2 mM KCl, 1 mM MgCl_2_, and 5 μM HEPES (pH 7.4). Afterwards, oocytes were washed with OR2 several times and then kept in OR2 solution supplemented with 1.8 mM CaCl_2_, 100 IU/ml penicillin, and 100 μg/ml streptomycin at 16°C.

To express exogenous proteins, cRNA was microinjected into oocytes using a nanoliter-injector system (Warner Instruments, Hamden, CT, USA). cRNA was *in vitro* transcribed from the linearized pGEMSH vectors which contained the cDNA clones for human TRPM3 (hTRPM3) (38) rat TRPM8, human M2 muscarinic (hM2) receptor, or GΔψ subunits by using the mMessage mMachine T7 Transcription Kit (Thermo Fisher Scientific). TRPM3 and TRPM8 mutants, that were used in this paper, were generated by the QuikChangeII XL Site-Directed Mutagenesis Kit from Agilent, and the mutated DNA constructs were confirmed by DNA sequencing. For co-expression of TRPM3 constructs and GΔ1ψ2 subunits, 40 ng of TRPM3 was coinjected with 5 ng Gβ1 and 5 ng Gγ2. In the case of co expressing TRPM3 and hM2 receptors, these two were injected at 1:1 ratio, 40 ng each. Oocytes were used for electrophysiological experiments after 48-72h incubation at 16°C.

### Two electrode voltage clamp experiments

Oocytes were placed in extracellular solution which contained 97 mM NaCl, 2 mM KCl, 1 mM MgCl_2_, and 5 μM HEPES, pH 7.4. Currents were measured with a protocol consisting a voltage step from the 0 mV holding potential to −100 mV, followed by a ramp to 100 mV once every 0.5 s with a GeneClamp 500B amplifier and analyzed with the pClamp 9.0 software (Molecular Devices). Currents were recorded by thin wall glass pipettes which contained inner filament and was filled with 1% agarose in 3 M KCl. In all TEVC experiments, different concentrations of PregS were applied to activate TRPM3 channels and various concentrations of menthol were used to trigger responses of TRPM8 channels. The hM2 receptor was activated by 5 μM of acetylcholine. For wortmannin experiments specifically, PregS or menthol-induced currents were measured, then the same oocyte was incubated with 35 μM wortmannin for 2h and currents were measured again using the same protocol.

### Excised inside-out patch clamp electrophysiology

Oocytes were placed in a recording chamber filled with bath solution which contained 97 mM KCl, 5 mM EGTA, 10 mM HEPES, pH 7.4. Before starting measurements, the vitelline layer was carefully removed with forceps without damaging the oocyte. Then a giga-ohm seal was formed using a borosilicate glass pipette (World Precision Instruments, Sarasota, Florida, USA) with resistance from 0.8 to 1 MΩ. The pipette was filled with a solution containing 97 mM NaCl, 2 mM KCl, 1 mM MgCl_2_, 5 mM HEPES, and 100 µM PregS at pH 7.4. Currents were measured by an Axopatch 200B amplifier and analyzed with the pClamp 9.0 software (Molecular Devices, Sunnyvale, CA, USA). Compounds were dissolved in the bath solution and were delivered to the inner side of cell membrane by a custom-made gravity driven perfusion system. Either 25 μM PI(4,5)P_2_, 25 μM PI(4)P or 10 μM AASt PI(4,5)P_2_ was applied in these experiments to reactivate TRPM3. At the end of every recording, 30 μg/ml Poly-Lys (Poly-K) was applied.

### Maintenance and Transfection of HEK293 cells

Human Embryonic Kidney 293 (HEK293) cells were purchased from American Type Culture Collection (ATCC, Manassas, VA, catalogue number is CRL-1573). HEK293 cells were cultured in the MEM supplemented with 10% FBS and 100 IU/ml penicillin plus 100 µg/ml streptomycin. Cells were incubated in 5% CO_2_ at 37°C. Cells were tested to confirm that they were not infected by the mycoplasma. Cells were used up to 25 passages and then discarded. Cells were transiently transfected with cDNA encoding different TRPM3 constructs (200-400ng) using the Effectene reagent (Qiagen). Mouse TRPM3α2 (mTRPM3α2) and its mutant were cloned into the bicistronic pCAGGS/ IRES-GFP vector. The components of rapamycin inducible pseudojanin phosphatases (40) were co transfected with mTRPM3α2 at 1:1 ratio.

### Whole cell patch clamp experiments

After 24h of transfection, cells were plated on poly-D-lysine coated 12mm cover slips. Experiments were performed 48-72 hours post transfection. Coverslips were placed in recording chamber filled with extracellular solution (137 NaCl, 5 KCl, 1 MgCl_2_, 10 HEPES and 10 glucose, pH 7.4). Since mouse TRPM3 constructs were in the background of bicistronic pCAGGS/ IRES-GFP vector and rapamycin inducible phosphatases were labeled with RFP, cells which showed both GFP and RFP fluorescence, were selected for the whole cell patch clamp experiments. Those patch pipettes were prepared from borosilicate glass capillaries (Sutter Instruments) using a P-97 pipette puller (Sutter Instrument) with a resistance of 2-4 MΩ. Those recording pipettes were filled with intracellular solution containing 140 mM potassium gluconate, 5 mM EGTA, 1 mM MgCl_2_, 10 mM HEPES, and 2 mM Na-ATP, pH 7.4. After formation of gigaohm-resistance seals, the whole cell configuration was established, and currents were recorded by applying a ramp protocol once every 1 s. The holding potential was 0 mV; followed by a −100 mV step for 100 ms; plus a ramp protocol from −100 mV to +100 mV over the period of 500 ms. All recording were made with an Axopatch 200B amplifier, filtered at 5 kHz, and digitized through Digidata 1440A interface. Data were collected and analyzed with the PClamp10.6 (Clampex) acquisition software (Molecular Devices, Sunnyvale, CA), and further analyzed and plotted with Prism 9 (Graphpad by Dotmatics). TRPM3 channels were activated by PregS and 100 nM of rapamycin was applied to activate phosphatases.

### Statistics

Statistical analysis was performed with Origin 2021 and Graphpad Prism 9. Data were plotted as mean ± SEM and scatter plots. Samples were not predetermined by any statistical method, however, they were similar to what is generally used in the field. All recordings were performed in random order. Statistical significance was evaluated with t-test, or analysis of variance with Bonferroni’s post hoc test, or the Kolmogorov Smirnov non-parametric test, as described in the figure legends. P values are reported in figures or figure legends.

## Computational Methods

### TRPM8 Experimental structure refinement and molecular docking of full-length PI(4,5)P_2_

The cryo-electron microscopy structure of full-length, apo TRPM8 from *Ficedula albicollis* (PDB-ID 6BPQ) (30), which contains several unresolved amino-acid ranges (∼4.1 Å resolution) as well as protein residues with missing atoms, was used as the starting configuration to generate a refined structural model of the TRPM8 channel. The Prime Loop Prediction (52) program and the Protein Preparation Wizard (53) (both distributed by Schrödinger, LLC, New York, NY, 2018) were used to perform the following tasks: (I) loop refinement by serial loop sampling, at the ultra-extended accuracy level. In particular, 4 unresolved amino-acid ranges in the transmembrane (TM) region were sampled, including 714 to 722, 819 to 822, 889 to 895 and 976 to 990 (sequence numbering as in *Ficedula albicollis*); (II) side chain prediction of protein residues with missing atoms, performed with no backbone sampling; (III) pKa prediction of protein residues at pH 7, followed by analysis and optimization of hydrogen-bond networks; (III) structure refinement via restrained minimization of heavy atoms (hydrogens not restrained) using the OPLS (54) force field. The minimization convergence criterion was set to 0.30 Å RMSD for heavy atom displacement. The resulting apo TRPM8 structure was then searched for putative ligand binding sites using SiteMap (35) (Schrödinger, LLC, New York, NY, 2018). Residues facing the topmost suitable site for ligand binding spot were used to define the docking space for putative PI(4,5)P_2_ binding modes. The program Glide (55) 1. Method and assessment of docking accuracy} (Schrödinger, LLC, New York, NY, 2018) was used to dock PI(4,5)P_2_ against TRPM8, using a rigid-receptor and flexible-ligand protocol. The ligand was prepared by using the default protocol of LigPrep (Schrödinger, LLC, New York, NY, 2018). Binding modes were ranked using the Glide standard precision (SP) scoring function. The best binding mode of PI(4,5)P_2_ against TRPM8 is shown in **Figure 2**. After our refined TRPM8-PI(4,5)P_2_ complex was generated and used for subsequent modeling of the TRPM3 channel as in a complex with PI(4,5)P_2_, seven additional experimental structures of TRPM8 became available (**Table S1**). Three of these structures report the TRPM8 channel in complex with PI(4,5)P_2_ as well as Ca^2+^ ions and/or small molecule ligands (28).

### TRPM3 Homology model and molecular docking of a truncated PI(4,5)P_2_ molecule

No experimental structure of the TRPM3 channel is currently available. The cryo-electron microscopy structure of the TRPM4 channel (3.1 Å resolution) in the apo state with short coiled coil from *Mus musculus* (PDB-ID 6BCJ) (29), was selected as the template to build a homology model of the human TRPM3 structure using the Swiss-Model Server (56) (https://swissmodel.expasy.org/), based on the human sequence UniProtKB: Q9HCF6. The choice of the template is exemplified in **Scheme S1**. Essentially, the closest relative to TRPM3 in the TRPM family (Cladogram) with an available structural template was selected, i.e. – TRPM4 (29). The cladogram was generated using Clustal Omega (https://www.ebi.ac.uk/), upon performing a multiple sequence alignment (default settings) (57). The Swiss-Model-generated protein structure of apo TRPM3 was then prepared for subsequent calculations using the Protein Preparation Wizard (53) (Schrödinger, LLC, New York, NY, 2018). Potential hot spots for PI(4,5)P_2_ binding to TRPM3 were defined by combining binding-site mapping results obtained using SiteMap (35) (Schrödinger, LLC, New York, NY, 2018) with sequence and structure alignments between the refined structural model of TRPM8 in complex with PI(4,5)P_2_ and the TRPM3 model (apo state) generated using Swiss-Model (56). Hence, the TRPM3 protein residues facing the most “druggable” binding spot were selected by homology and used to center the docking grid for subsequent docking of PI(4,5)P_2_. The best binding mode of a truncated version of PI(4,5)P_2_ against TRPM3 is shown in **Figure 1**. As a matter of fact, due to the extreme flexibility of the lipid tail, the docking algorithm failed in generating binding poses for the full length PI(4,5)P_2_ lipid. Instead, starting from the PI(4,5)P_2_ head group, a series of truncated versions of a growing lipid were docked successfully against the binding site on TRPM3 until a maximum tail length was reached (our truncated lipid is similar to the syntethic diC_8_ PI(4,5)P_2_ molecule is experimentally functional in activating TRPM3 (21)). For simplicity, in this work the PI(4,5)P_2_ lipid with truncated tails, which was modeled in complex with TRPM3, is referred to as PI(4,5)P_2_. Note that in for TRPM8, the PI(4,5)P_2_ molecule was modeled as a full-length lipid. As for the molecular docking, we used the same protocol implemented for TRPM8. Related figures were generated using the Visual Molecular Dynamics (VMD) molecular visualization program (58).

### Comparisons of TRPM8 and TRPM3 models

Structural superimpositions of the model of TRPM8 in complex with (full length) PI(4,5)P_2_ and that of TRPM3 in complex with (truncated) PI(4,5)P_2_ were performed upon structural superposition based on sequence alignments (using the algorithm Needelman-Wunsh with BLOSUM-62 matrix). Alignments were generated using the Match Maker tool in UCSF Chimera (59), version 1.15, and analyzed in VMD. The following experimental structures were aligned to the TRPM8/PI(4,5)P_2_ model: the TRPM8 in complex with the menthol analog WS-12 and PI(4,5)P_2_ (PDB-ID: 6NR2), and two crystal complexes of TRPM8 with icilin (PDB-ID: 6NR3 and 6NR4), PI(4,5)P_2_, and calcium (28). The following structures were aligned to the TRPM3/PI(4,5)P_2_ model: the experimental structure of TRPM4 (29) and TRPM3 models from AlphaFold (31,32). At the time of writing, 4 AlphaFold models were available of TRPM3, each from a different organism (UniProt sequence-id: Q9HCF6 (human; Fig. S1), J9S314 (*Mus musculus*), F1QYX6 (Danio rerio) and F1LN45 (Rattus norvegicus). All 4 structures were superimposed (not shown), revealing striking structural similarities. Related figures were generated using VMD.

### ΔΔG calculations

Changes in the binding affinity (or Gibbs free energy of binding, ΔΔ*G* in kcal/mol) of PI(4,5)P_2_ to TRPM3 were calculated upon mutating key binding residues in the putative PI(4,5)P_2_ binding site. To do so, a computational protocol previously used with similar systems was employed (25,60). Essentially, residue mutations and ΔΔ*G* calculations were performed on the generated TRPM3 model bound to a truncated PI(4,5)P_2_ molecule, i.e. – the native structural complex. The “Residue-Scanning and Mutation” tool from BioLuminate (61) (Schrödinger, LLC, New York, NY, 2018), was used to perform calculations upon mutating residues listed in **Table S2**. These residues were also mutated experimentally. The structural complexes were then refined by side-chain prediction with backbone sampling of the mutated residue, before a minimization in the region around the mutation site was performed to relax the and optimize the side-chain interactions with the bound lipid. Systems were prepared for the calculations using the Protein Preparation Wizard (53) (Schrödinger, LLC, New York, NY, 2018). Affinity changes were plotted using Microsoft Excel (https://www.microsoft.com/). Related figures were generated using Maestro (Schrödinger, LLC, New York, NY, 2018).

## DATA AVAILABILTY

All data are contained in the manuscript and in the supporting information. Computational data will be made available for academic use upon request to the authors.

## ACKNOWLEDGMENTS

This study was supported by grants NSNS055159 to TR and GM131048 to VC and TR. The hTRPM3_1325_ clone in a mammalian expression vector was provided by C Harteneck (Eberhard Karls University Tubingen, Tubingen, Germany). The mTRPM3α2 clone was a kind gift from Drs. Veit Flockerzi and Stephan Phillipp, and the rat TRPM8 clone was provided by Dr. David Julius. The human muscarinic M2 receptor, and the Gβ1 and Gγ2 clones were provided by Dr. Diomedes Logothetis. This research includes calculations carried out on HPC resources at Temple University supported in part by the National Science Foundation through major research instrumentation grant number 1625061 and by the US Army Research Laboratory under contract number W911NF-16-2-0189. Molecular graphics and analyses performed with UCSF Chimera, developed by the Resource for Biocomputing, Visualization, and Informatics at the University of California, San Francisco, with support from NIH P41-GM103311.

## CONFLICT OF INTEREST

The authors declare that they have no conflicts of interest with the contents of this article.

---

**Table S1.** Experimental structures of TRPM8 available (https://www.rcsb.org). The template selected for generating the refined model of TRPM8 in complex with PI(4,5)P_2_ is 6BPQ. This is the only TRPM8 structure released before the TRPM3 model was built (the latter was built on TRPM4 6BCJ as the template).

| PDB-ID | Channel | Release | State |
| --- | --- | --- | --- |
| 6BPQ | TRPM8 | Before TRPM3 model | Ligand free |
| 6NR2 | TRPM8 | After TRPM3 model | Menthol analog WS-12 and PI(4,5)P<sub>2</sub> |
| 6NR3 | TRPM8 | After TRPM3 model | Icilin, PI(4,5)P<sub>2</sub>, and Ca<sup>2+</sup> (high occupancy) |
| 6NR4 | TRPM8 | After TRPM3 model | Icilin, PI(4,5)P<sub>2</sub>, and Ca<sup>2+</sup> (low occupancy) |
| 6O6A | TRPM8 | After TRPM3 model | Ligand free |
| 6O6R | TRPM8 | After TRPM3 model | AMTB-bound state |
| 6O72 | TRPM8 | After TRPM3 model | TC-I 2014-bound state |
| 6O77 | TRPM8 | After TRPM3 model | Calcium-bound state |

**Table S2.** Change in the binding affinity (Δ affinity or ΔΔG) for PI(4,5)P_2_ to the human TRPM3 model (Q9HCF6) predicted upon mutating key residues responsible for binding. Sequence notation for both the experimental and the computational isoforms are reported, as well as the corresponding residues mapped on the TRPM8 binding site (*Ficedula albicollis)*.

| TRPM3 Mutant (Exp. Isoform) | TRPM3 Mutant (Comp. Isoform) | $\Delta$ Affinity (kcal/mol) | Refined TRPM8 Model (on 6BPQ) |
| --- | --- | --- | --- |
| N991A | N1016A | 7.90 | S849 |
| K992A | K1017A | 16.4 | R850 |
| R1131Q | R1156Q | 8.60 | R997 |
| W761F | W786F | 6.00 | W682 |
| R1141Q | R1166Q | 1.10 | R1007 |
| K1128Q | K1153Q | 1.20 | K994 |

**Figure S1.**
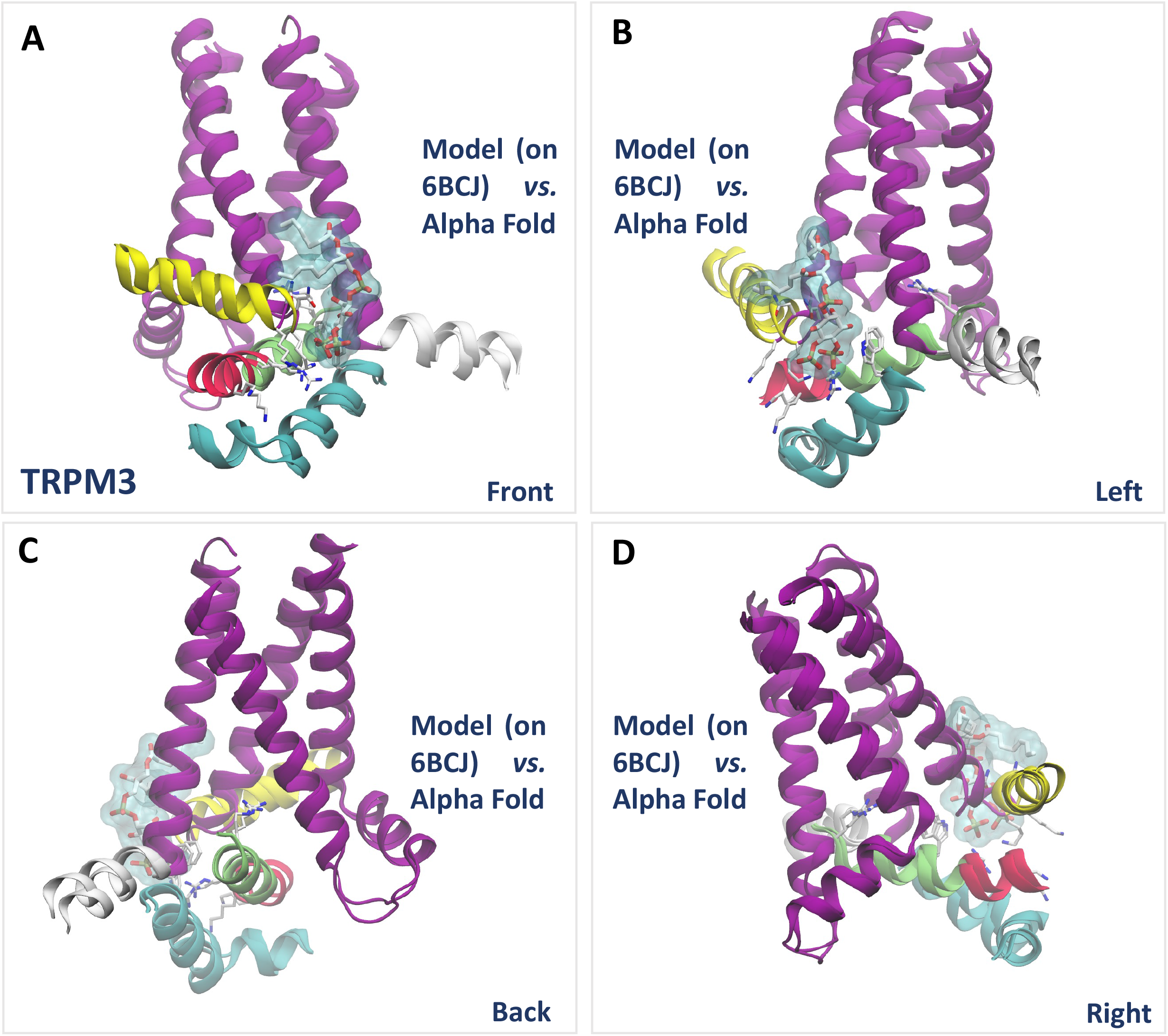
Structural comparisons of the TRPM3 model in complex with PI(4,5)P_2_ with a ligand-free model from AlphaFold. (**A to D**) Close-up views of structural superposition of the PI(4,5)P_2_ binding site in two different TRPM3 structures, namely the TRPM3 model (built on TRPM4, 6BCJ) (29) in complex with PI(4,5)P_2,_ which was built in this work, and the ligand-free protomer of human TRPM3 from AlphaFold (Q9HCF6) (31). All representations are reproduced as in Figure 1.

**Figure S2.**
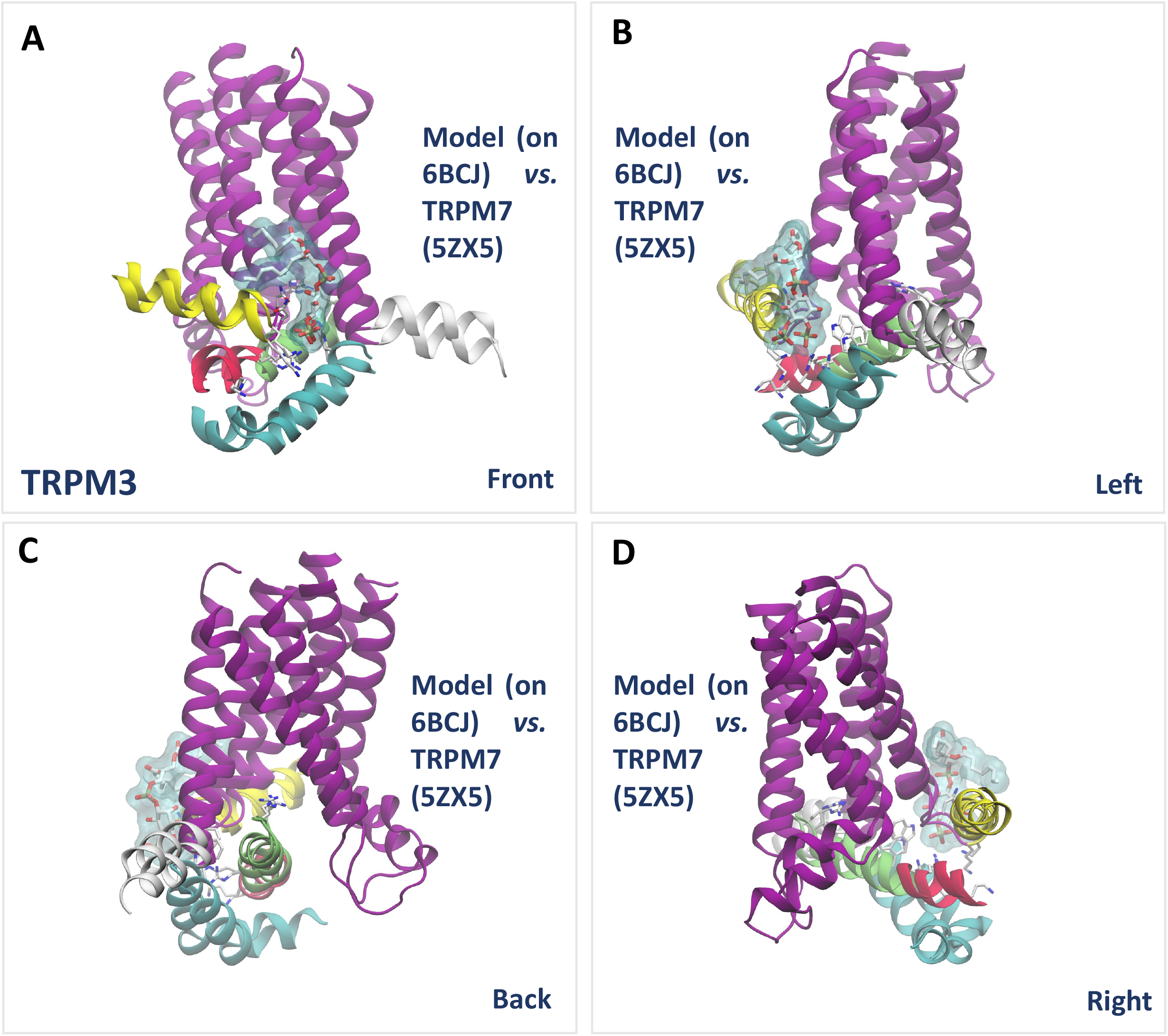
Structural comparison of TRPM3 models. (**A to D**) Close-up views of structural superposition of the PI(4,5)P_2_ binding site in two different TRPM3 models (both built in this work). The first is the main TRPM3 model in complex with PI(4,5)P_2,_ built using TRPM4 (6BCJ) (29) as the template. The second is a model of ligand-free TRPM3 obtained using the cryo-EM structure of TRPM7 in EDTA (5XZ5) (33) as the template. All representations are reproduced as in Figure 1.

**Figure S3.**
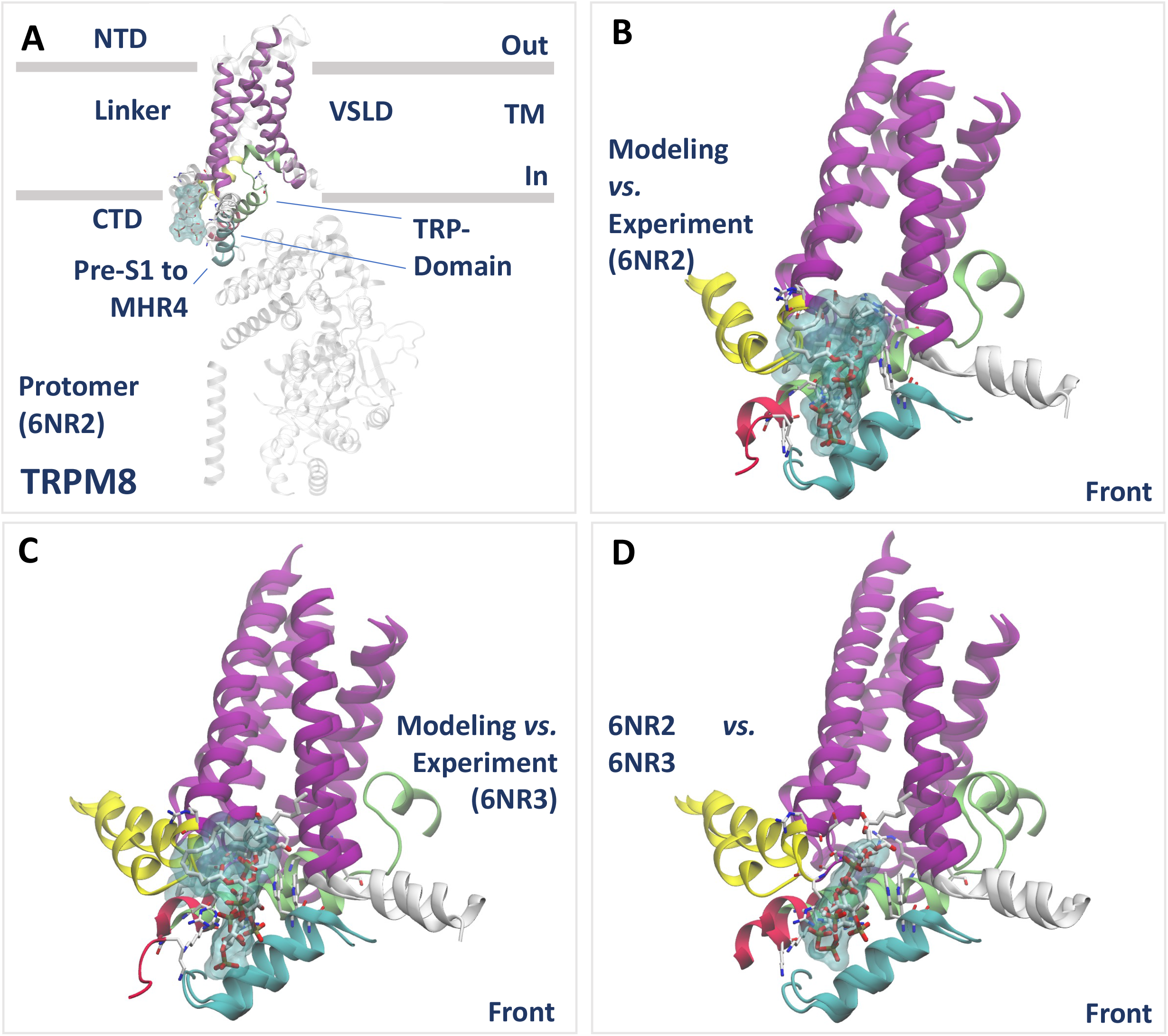
Structural comparisons of the model of TRPM8 in complex with PI(4,5)P_2_ with experimental structures of TRPM8. (**A**) Protomer view from the transmembrane (TM) plane of the cryo-EM structure of TRPM8 in complex with the menthol analog WS-12 and PI(4,5)P_2_ (6NR2) (28). The structure compares with the refined model of TRPM8 in complex with PI(4,5)P_2_ (Figure 2). (**B to D**) Close-up views of structural superposition of the PI(4,5)P_2_ binding site in different TRPM8 structures. In (B), the refined model of TRPM8 in complex with PI(4,5)P_2_ is superposed to the cryo-EM structure shown in (A). In (C), the model is superposed to the cryo-EM protomer of TRPM8 in complex with icilin, PI(4,5)P_2_ and calcium (6NR3) (28). In D, protomers from the two cryo-EM structures of TRPM8 in complex with PI(4,5)P_2_ are superposed. All representations are reproduced as in Figure 1.

**Figure S4.**
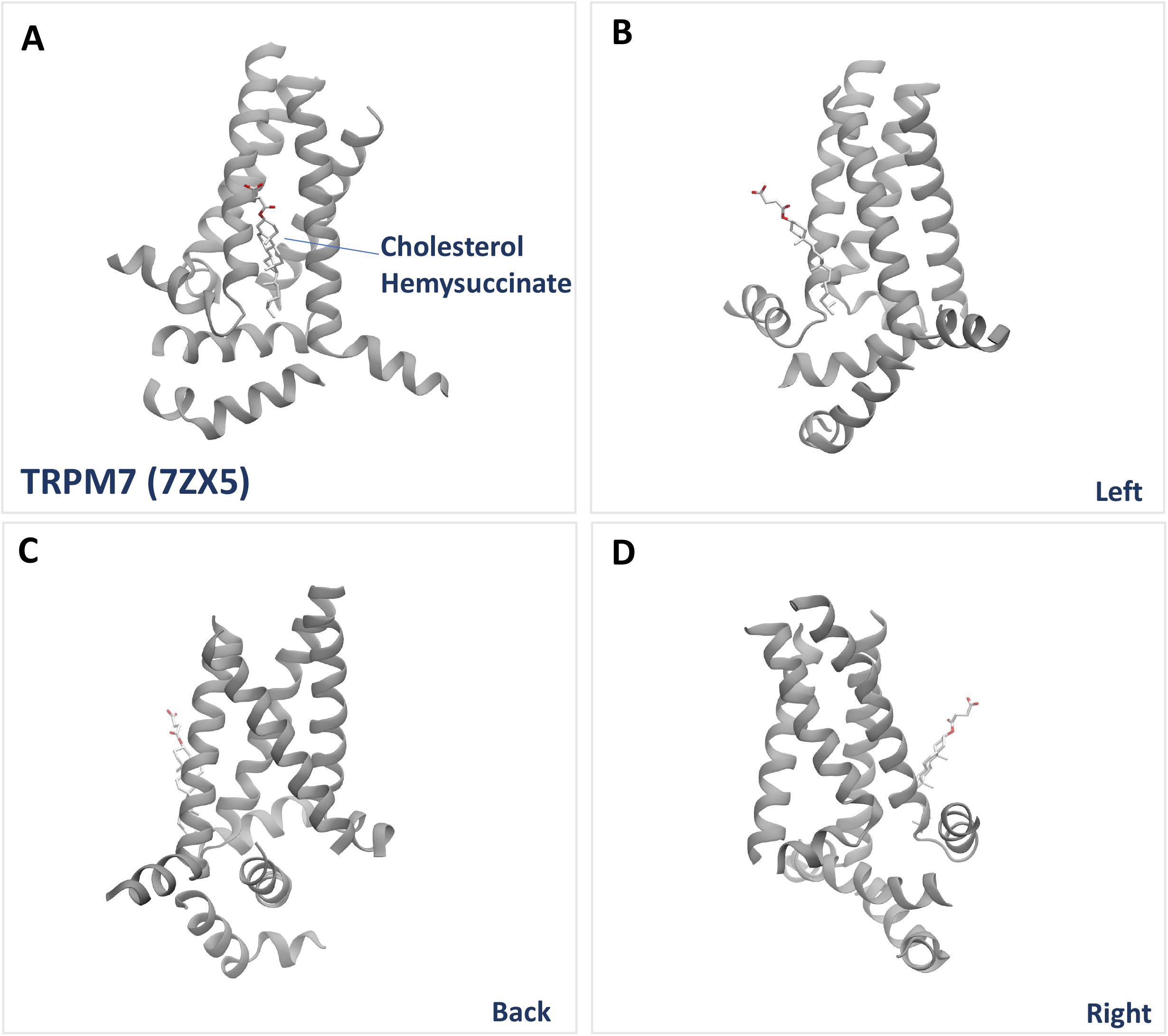
Lipid binding site in the experimental structure of TRPM7. (**A to D**) Close-up views of a putative lipid binding site in the cryo-EM structure of TRPM7 (5XZ5) (33). The site superposes well with the PI(4,5)P_2_ binding site identified in our TRPM3 model (Figure 1E), and in our TRPM8 model (Figure 2E) and in the cryo-EM structure of TRPM8 bound to PI(4,5)P_2_ (**Figure S3**). In the cryo-EM structure of TRPM7, a molecule of detergent cholesterol-hemisuccinate (CHS) is bound to TRPM7 at the putative lipid binding site. Protein atoms are shown in new cartoon representation, colored in grey. CHS atoms are shown in licorice representation, with C, N, O atoms colored in white, blue and red, respectively.

**Figure S5.**
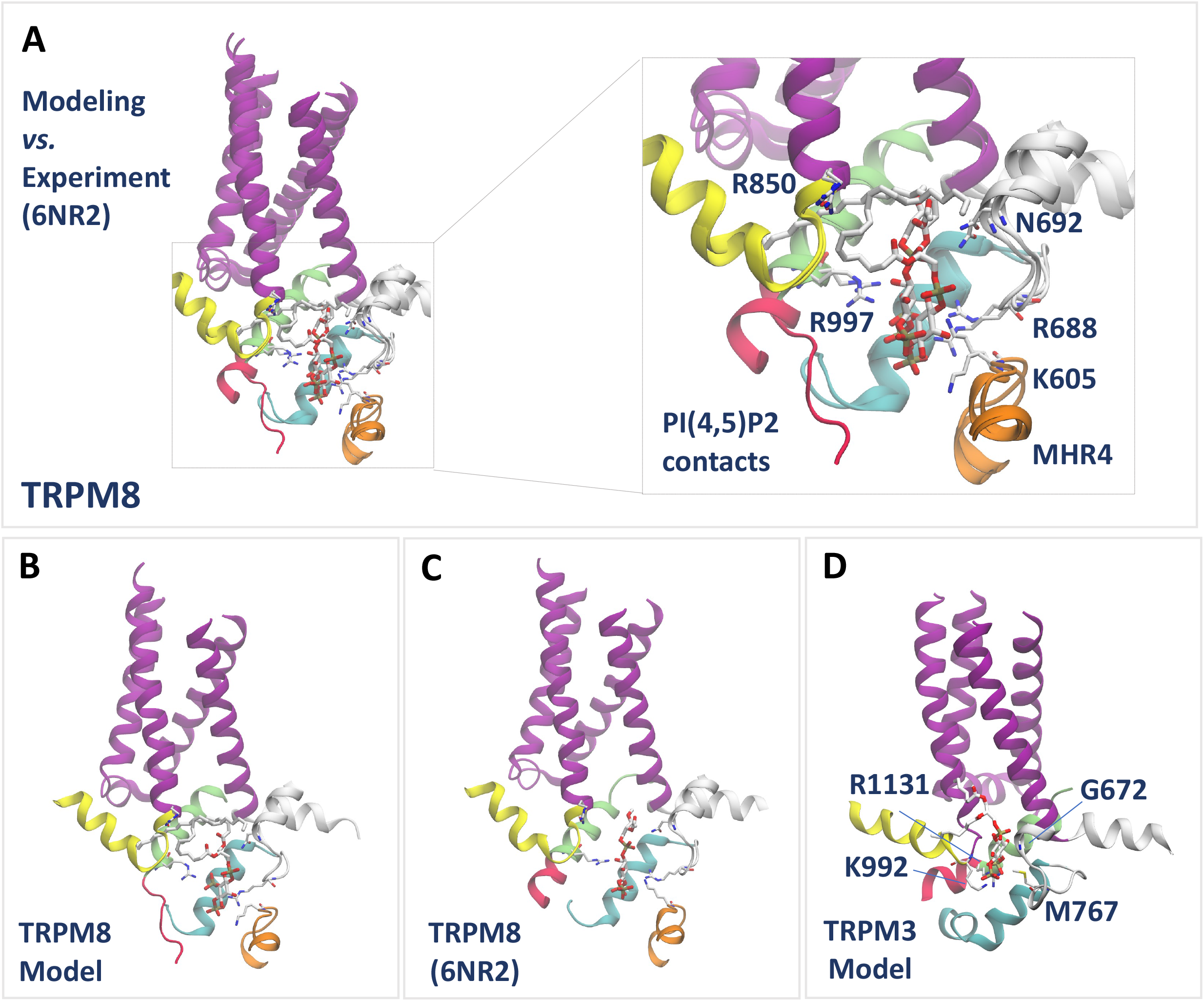
Structural comparisons of the PI(4,5)P_2_ binding site in TRPM3 and TRPM8 structures. **(A)** Close-up view with relative zoom-in (inset) of the PI(4,5)P_2_ binding site in the refined model of TRPM8 in complex with PI(4,5)P_2_ (Figure 2) superposed to the cryo-EM structure of TRPM8 in complex with the menthol analog WS-12 and PI(4,5)P_2_ (6NR2) (28). In the experimental structure, both WS-12 and PI(4,5)P_2_ are not shown. The experimentally determined PI(4,5)P_2_ contact residues (28) are shown in the inset. (**B to D**) Close-up views of (**B**) the refined model of TRPM8 in complex with PI(4,5)P_2_; (**C**) the structure of TRPM8 in complex with the menthol analog WS-12 and PI(4,5)P_2_ (6NR2); and (**D**) the TRPM3 model (Figure 1) in complex with PI(4,5)P_2_. The TRPM3 residues equivalent to the experimentally determined PI(4,5)P_2_ contacts (28) are shown. Comparisons of the PI(4,5)P_2_ site between TRPM8 and TRPM3 indicate striking structural similarities, with the exception of the MHR4 region (in orange new-cartoon representation), which is not conserved in TRPM3. All representations are reproduced as in Figure 1.

**Figure S6.**
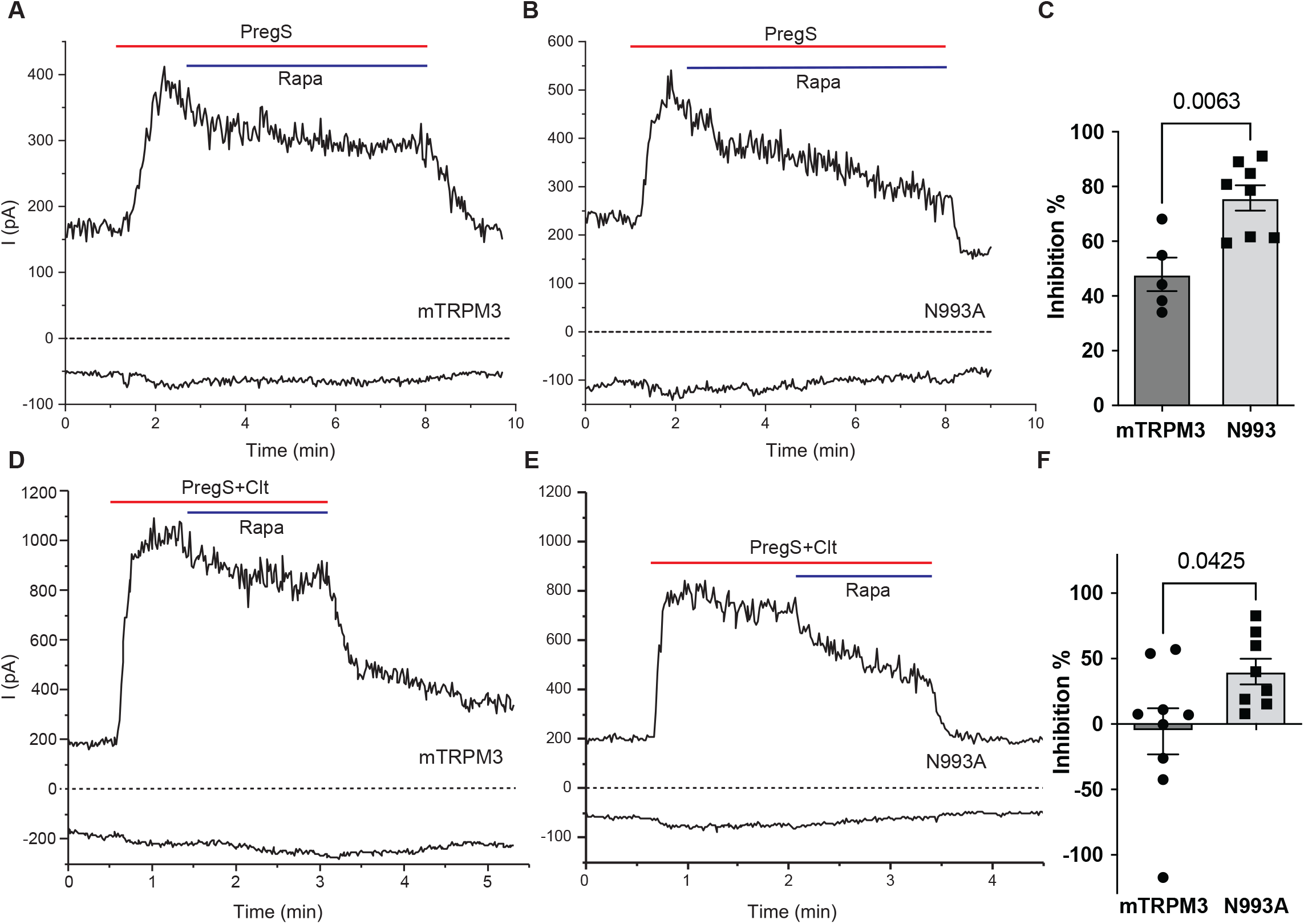
Mutating putative PI(4,5)P_2_ interacting residues increases sensitivity of TRPM3 to inhibition by PI(4,5)P_2_ depletion in HEK cells. HEK293 cells were transiently transfected with mouse TRPM3α2 (mTRPM3α2) or its mutant N993A, and the rapamycin inducible pseudojanin phosphatases constructs. Whole cell patch clamp electrophysiology was performed by using ramp protocols from - 100 mv to 100 mv as described in the Methods section. (A-B) Representative traces of mTRPM3α2 (A) and N993A (B). Applications of 25 μM PregS and co application of 100 nM rapamycin are indicated by red and blue lines respectively. Top traces show currents at +100 mV; dash lines indicate zero current; bottom traces show currents at −100 mV. (C) Data summary of the percentage of inhibition caused by PI(4,5)P_2_ depletion (rapamycin). Decreased Current amplitudes after rapamycin were normalized to PregS induced current amplitudes. (D-E) Representative traces of mTRPM3α2 (D) and N993A (E). Applications of 25 μM PregS and 10 μM Clotrimazole (Clt) are indicated by red lines. Applications of 100 nM rapamycin are indicated by blue lines. (F) Data summary of the percentage of inhibition caused by PI(4,5)P_2_ depletion. Decreased Current amplitudes after rapamycin were normalized to co application of PregS and clotrimazole (Clt) induced current amplitudes. Statistical significance was calculated by T test. Bar graphs show mean ± SEM and scatter plots, representing individual measurements from three independent transfections.

**Figure S7.**
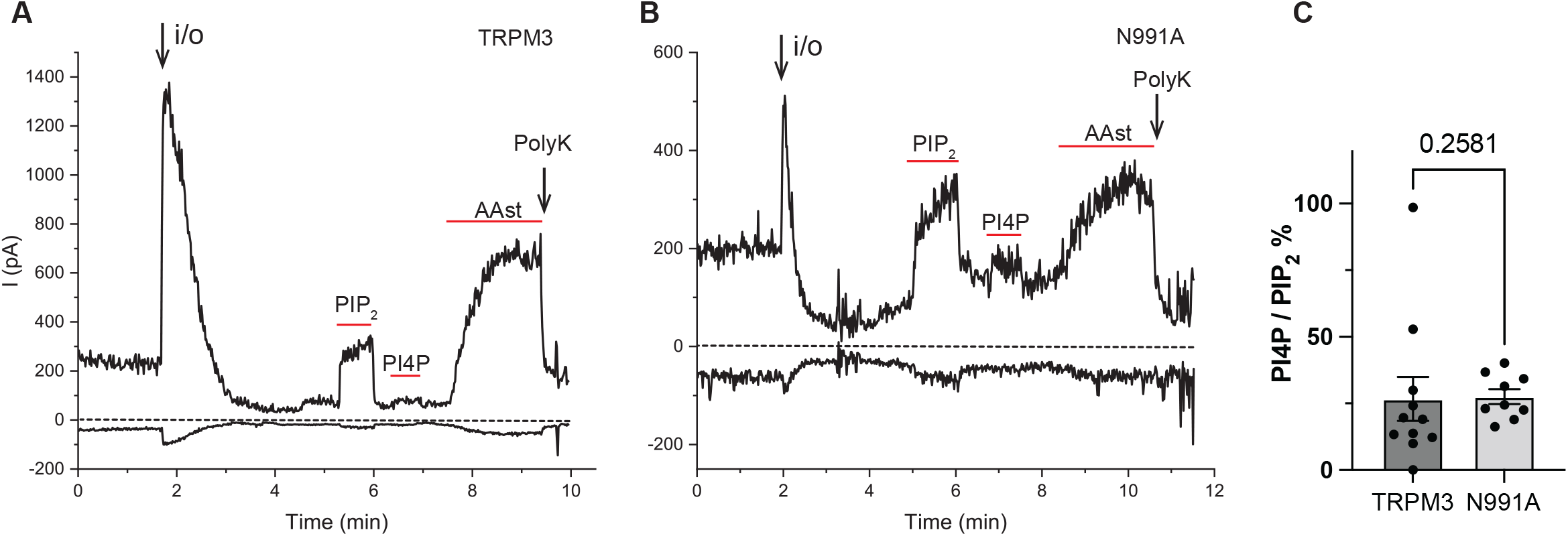
Mutating putative PI(4,5)P_2_ interacting residues does not change the relative effect of PI(4)P compared to PI(4,5)P_2_. hTRPM3 or its mutant N991A was expressed in oocytes and excised inside-out patch clamp electrophysiology was performed using ramp protocol from −100 mV to 100 mV as described in the Methods section. (A-B) Representative traces of hTRPM3 (A) and N991A (B). Top traces show currents at +100 mV; dash lines indicate zero current; bottom traces show currents at −100 mV. The formation of inside out configuration (i/o) is indicated by the arrow. Applications of 25 μM diC_8_ PI(4)P, 25 μM diC_8_ PI(4,5)P_2_ and 10 μM AAst PI(4,5)P_2_ are indicated by red lines, 30 μg/ml Poly-Lys (Poly K) was applied at the end and indicated by the second arrow. The patch pipettes contained 100 μM PregS to activate TRPM3 channels. (C) Data summary of the relative effect of PI(4)P compared to PI(4,5)P_2_. Current amplitudes induced by PI(4)P was normalized to current amplitudes induced by PI(4,5)P_2_. Kolmogorov-Smirnov non-parametric test was used to analyze data and P value was reported on the bar graph.

**Figure S8.**
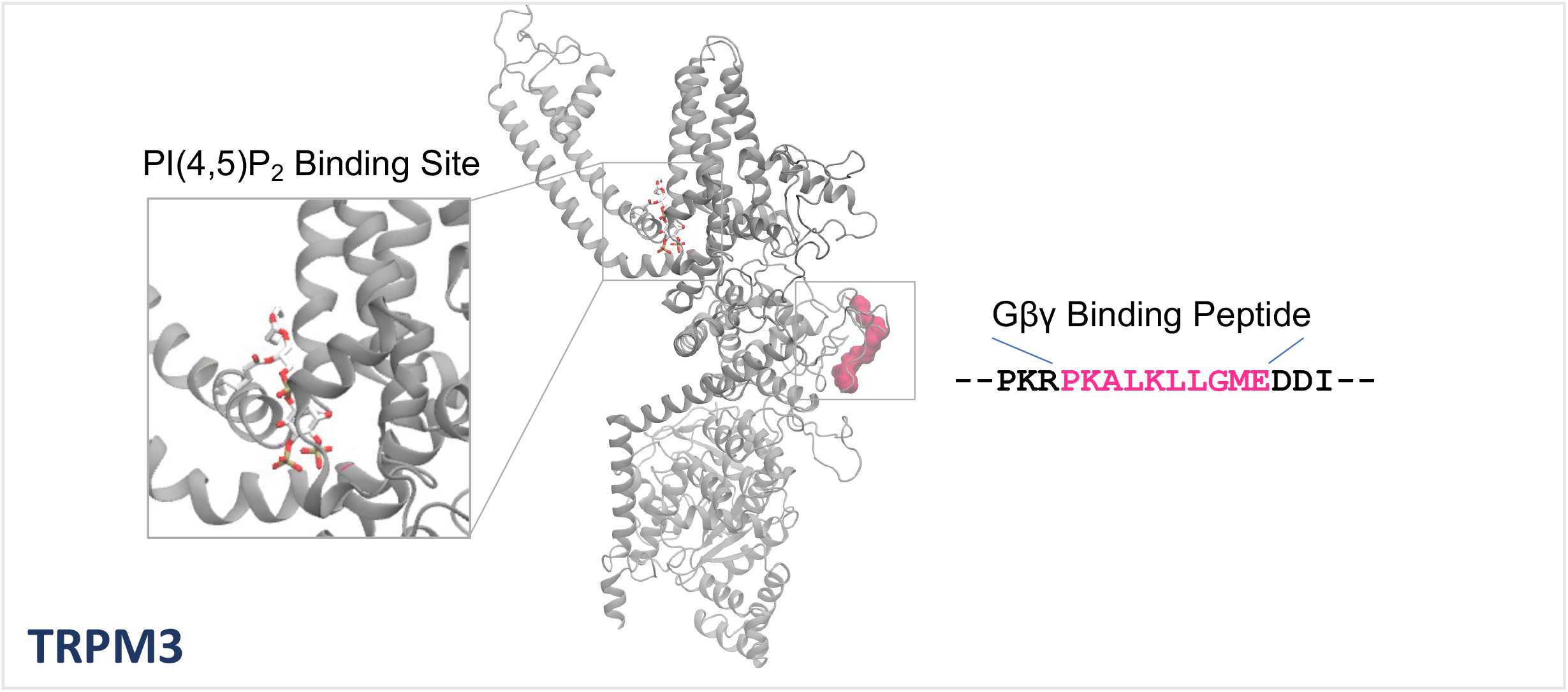
Location of the Gβγ binding peptide on TRPM3. (**Central**) View of a protomer of the TRPM3 model in complex with PI(4,5)P_2_, highlighting the location of the Gβγ-binding peptide (bright red) (12). (**Left inset**) A zoom-in view of the PI(4,5)P_2_ binding site is shown. (**Right**) The Gβγ-binding peptide sequence (in TRPM3) is shown (bright red). Protein atoms are shown in new cartoon representation, colored in grey. The Gβγ-binding region is shown in surface representation (bright red). PI(4,5)P_2_ atoms are shown in licorice representation, with C, N, O atoms colored in white, blue and red, respectively.

## Notes

### Competing Interest Statement

The authors have declared no competing interest.

